# Actions of the Anti-Seizure Drug Carbamazepine in the Thalamic Reticular Nucleus: Potential Mechanism of Aggravating Absence Seizures

**DOI:** 10.1101/2025.02.03.636080

**Authors:** Sung-Soo Jang, Nicole Agranonik, John R. Huguenard

**Author notes:** Corresponding author: John R Huguenard, Stanford Neuroscience Building S189, 290 Jane Stanford Way, Stanford, CA 94305-5088.

## Abstract

Carbamazepine (CBZ) is a widely used antiepileptic drug effective in managing partial and generalized tonic-clonic seizures. Despite its established therapeutic efficacy, CBZ has been reported to worsen seizures in another form of epilepsy, generalized absence seizures, in both clinical and experimental settings. In this study, we focused on thalamic reticular (RT) neurons, which regulate thalamocortical network activity in absence seizures, to investigate whether CBZ alters their excitability, thereby contributing to the exacerbation of seizures. Using ex vivo whole-cell patch-clamp electrophysiology, we found that CBZ selectively inhibits the tonic firing of RT neurons in a dose-dependent manner without affecting burst firing. At the RT-thalamocortical (RT-TC) synapse, CBZ significantly increases the failure rate of GABAergic synaptic transmission, with greater effects on somatostatin (SST) – than parvalbumin (PV) - expressing RT neurons. In vivo EEG recordings and open-field behavior in Scn8a^med+/-^ mouse model confirmed that CBZ treatment exacerbates absence seizures, increasing both seizure frequency and duration while reducing locomotor activity. In addition, CBZ further amplifies the pre-existing reduction in tonic firing of RT in Scn8a^med+/-^ mice. These findings uncover a novel mechanism by which CBZ exacerbates absence seizures through selective inhibition of RT neuron excitability and disruption of GABAergic synaptic transmission. This work provides mechanistic insights into the paradoxical effects of CBZ and suggest potential avenues for optimizing epilepsy treatment strategies.

**Scientific Significance:** This study addresses the clinical paradox in which CBZ, a widely prescribed antiepileptic drug, paradoxically aggravates absence seizures. Understanding the cellular mechanisms behind this phenomenon is critical for improving epilepsy treatments. Here, using electrophysiology recordings from intact thalamocortical slices and SCN8a^med+/-^ mice, an absence seizure animal model, we demonstrate that CBZ selectively inhibits tonic firing of RT neurons and their output to thalamocortical circuits, with a more pronounced effect in SCN8a^med+/-^ mice. These novel findings provide a mechanistic explanation for CBZ’s paradoxical aggravation of absence seizures, offering a framework for understanding the pharmacological effects of other anti-epilepsy drugs and guiding the development of more effective therapeutic strategies for epilepsy.

## Introduction

Carbamazepine (CBZ) is widely prescribed for epilepsy management, effectively treating partial seizures, generalized tonic-clonic seizures, and mixed seizure patterns. Its primary mechanism of action is proposed to be suppression of repetitive neuronal firing through use-dependent blockade of voltage-gated sodium channels, thereby reducing seizure-related neuronal hyperactivity. However, clinical evidence indicates that CBZ exacerbates absence seizures (1–4) characterized by transient episodes of impaired consciousness and distinctive 3 Hz spike-and-wave discharge (SWD) on electroencephalography (EEG) in patients with generalized epilepsies. Studies in the Generalized Absence Epilepsy Rats from Strasbourg (GAERS) have documented that CBZ aggravates 3-6 Hz SWD through GABAa receptor-dependent mechanisms in the thalamic ventrobasal complex (VB) (5). In addition, CBZ has been shown to augment spontaneous SWDs in the stargazer mutant mouse model of absence seizures (6).

The thalamic reticular (RT) nucleus is a specialized structure composed of GABAergic neurons, serving as a major source of inhibition to thalamocortical (TC) neurons (7–9). Selective activation of RT neurons through optogenetic stimulation induces thalamocortical neuron burst firing and rhythmic activity, hallmark features of absence seizure (10). RT neurons are predominantly parvalbumin-expressing (PV), although recent studies have identified a subpopulation of somatostatin-positive (SOM) neurons within this nucleus (11). Notably, PV and SOM neurons differentially regulate distinct thalamocortical circuits. SOM-expressing RT neurons are implicated in the modulation of gamma rhythms and the processing of visual information in the primary visual cortex (V1) (12), whereas PV-expressing RT neurons contribute to rhythmic activity that underlies somatosensory behavior and the generation of absence seizures (11, 13, 14). Disruption of PV-expressing RT neuron activity is sufficient to induce spike-and-wave discharges (SWD) on an electroencephalogram (EEG) (13), highlighting their critical role in the pathophysiology of absence seizure. Hemizygous loss of *SCN8a*, which encodes the voltage-gated sodium channel Nav1.6, leads to reduced tonic firing of RT neurons and impaired intra-RT synaptic connectivity, contributing to thalamocortical network hypersynchrony and spontaneous absence seizures (15). Considering the central role of RT neurons in the modulation of absence seizures and the widespread use of CBZ in epilepsy treatment, it is to understand whether CBZ influences electrophysiological properties within RT neurons, leading to the aggravation of absence seizure.

In this study, we show the following: (1) at clinically relevant concentrations (30 µM), CBZ selectively inhibits tonic firing without affecting burst firing of RT neurons; (2) CBZ increases the failure rate of GABAergic transmission at the RT-TC synapse, with greater effects observed in SOM- vs PV-expressing RT neurons; (3) In SCN8a^med+/^ mice, which exhibit spontaneous absence seizures, CBZ increases both seizure frequency and duration while inducing hypo-locomotor activity; (4) CBZ inhibits RT excitability more prominently in SCN8a^med+/-^ mice compared to SCN8a^WT^ mice. These novel findings enhance our understanding of the mechanisms underlying the paradoxical effect of CBZ in absence seizure, providing valuable insights for refining the therapeutic management of epilepsy.

## Results

### CBZ selectively inhibits Tonic mode of RT firings in an activity-dependent manner

Given that CBZ has been shown to suppress sustained neuronal firing in a variety of cell types (16–19), we hypothesized that CBZ would similar effects on RT neurons. To test this hypothesis, we performed whole-cell patch clamp recordings on RT neurons using acute horizontal slices that preserved an intact thalamocortical circuit (Fig. 1A). RT neurons exhibited three distinct firing patterns: burst firing, burst followed by tonic firing, and tonic firing, elicited by weak, moderate, and strong membrane depolarization, respectively (Fig.1B). We first examined the dose-dependent effects of CBZ on RT neuronal firing at concentrations of 20, 30, and 50 µM CBZ, which are within the therapeutic reference range (4 - 12 ug/ml) for humans (20, 21). Current-clamp recordings were performed using current steps from 0 to 210 pA for 1 second. CBZ inhibited action potential (AP) firing in a dose-dependent manner, which was fully recovered after a 15-minute washout (SI Appendix, Fig.S1 A-D). We confirmed recovery in all findings, but for clarity these washout results are not included in all figures. To further explore the mechanisms underlying CBZ’s effects, we expanded the protocol to include depolarizing currents from 0 to 300 pA for 1 s (Fig. 1C). Given that 50 μM is near the top of the clinically relevant range (20), we selected a concentration of 30 μM (19), an effective dose with minimal evidence of side effects for the subsequent experiments in this study. Upon incubation with 30 μM CBZ for 10-minutes, we analyzed AP firings across three distinct time intervals within the spike train: early (0 - 100 ms, a period dominated by early burst firing), middle (500 - 600 ms, a period of transition between burst and tonic firing), and late (900 - 1000 ms, a period restricted to only tonic firing). AP numbers were significantly reduced by CBZ only during the middle and late time periods, while AP firing during the early period remained unaffected (Fig. 1D). We next investigated the effects of CBZ on fundamental AP properties including overshoot (mV), fast AHP (mV), Half-width (ms), AP threshold (mV), and AP amplitude (mV) in APs from the first 3 spikes and the last 3 spikes of each spike train. Comparisons between the first 3 and last 3 spikes in the spike train revealed significant CBZ-induced alterations in the AP properties, especially of the last 3 spikes (Fig. S2). Phase-plane analysis of the somatic AP waveform further supported these findings, demonstrating changes in AP rise and fall rates (Max dV/dt, Min dV/dt) and axon initial segment (AIS)-to-soma propagation (1st and second d^2^V/dt^2^ peaks) for the last 3 but not the first 3 spikes (Fig. 1E). Based on these observations, we hypothesized that CBZ selectively targets late tonic firing through a use-dependent block mechanism while sparing early burst firing. To test directly for an effect on tonic firing, we held the membrane potential at – 60 mV to inactivate T-type calcium channels, minimizing their contribution to spike generation, and thus reducing burst firing. Under these conditions, CBZ inhibited AP firing across all current steps (30 – 300 pA) and time intervals (0 - 100, 500 - 600, and 900 - 1000 ms) (Fig. S3B and 3C). Additionally, CBZ affected key AP properties, including overshoot, halfwidth, and AP amplitude (Fig. S3D). Collectively, these findings highlight that CBZ selectively modulates RT neuronal excitability, with pronounced effects on tonic firing patterns, especially with depolarized resting membrane potentials.

**Figure 1.**
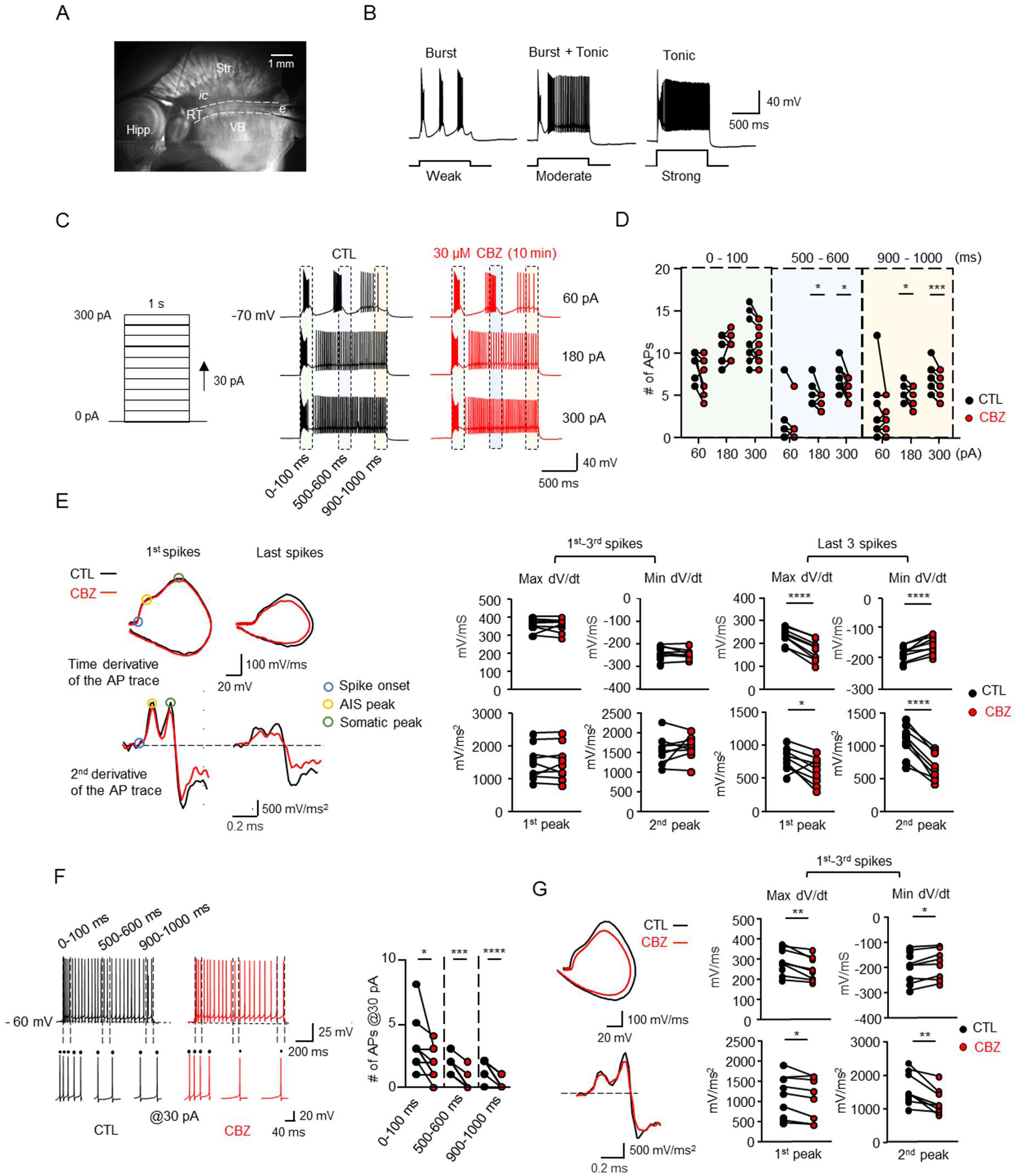
Selective Inhibition of Tonic Firing in RT Neurons by CBZ. (A) A microscopic image showing RT region and adjacent ventrobasal complex (VB) in a horizontal mouse brain slice (270 µm thickness). (B) Representative traces of action potentials (APs) illustrate two distinct firing modes in RT neurons: burst firing and tonic firing, elicited by weak, moderate, and strong depolarizing current injections, respectively. (C) Stimulation protocol (left) and corresponding AP traces (right) in RT neurons during a 1-second stimulation with increasing current injections (0–300 pA in 30 pA increments) before (black) and after (red) a 10-minute incubation with 30 µM CBZ. Cyan, gray, and yellow shaded areas highlight the first 100 ms, the 500–600 ms, and the final 100 ms of the stimulation epoch, respectively. (D) Quantification of the number of APs at specified time intervals (0–100 ms, 500–600 ms, 900–1000 ms) in response to current injections of 60, 180, and 300 pA, as color-coded in (C). Individual cells are represented by filled circles (n = 9). (E) Representative traces (left) showing the first derivative (dV/dt) and second derivative (d^2^V/dt^2^) waveforms of the first spike and last spike recorded at 300 pA. The graphs (right) compare the average maximum and minimum dV/dt, as well as the first and second peaks of d^2^V/dt^2^, between the first three and last three APs at 300 pA, before (black) and after (red) CBZ incubation (n = 9). (F) Representative traces (left) and quantification (right) of the number of spikes during tonic firing induced by a 30 pA current injection from a holding potential of –60 mV, over a 1-second epoch, before (black) and after (red) 10-minute CBZ incubation (n = 9). Individual spikes are represented by black dots. (G) Representative traces (left) showing the first derivative dV/dt (top) and the second derivative d^2^V/dt^2^ (bottom) waveforms for the first and last APs during tonic firing at 30 pA. Graph (right) compares average dV/dt, d^2^V/dt^2^, and their respective peaks between the first three and last three APs at 30 pA before (black) and after (red) CBZ incubation. Paired t-test was used for statistical analysis. *p < 0.05, **p < 0.01, ***p < 0.001, ****p < 0.0001.

### CBZ disrupts GABAergic transmission at the RT-TC synapse

To evaluate the effect of CBZ on synaptic output at the RT to TC synapse, we utilized VGAT-ChR2 mice (VGAT-mhChR2-YFP), in which GABAergic neurons, including RT cells, express channelrhodopsin- 2 (ChR2) coupled with enhanced yellow fluorescent protein (hChR2-eYFP). Optogenetic stimulation using a 470 nm LED light reliably evoked postsynaptic currents (PSCs) in TC neurons under conditions where glutamatergic transmission was blocked by 1 mM kynurenic acid (Fig. 2A). To investigate how CBZ affects GABAergic synaptic transmission at RT to TC synapses, we applied 470 nm LED stimulation at frequencies of 5, 10, 20, 30, and 50 Hz for 5 seconds. CBZ significantly increases the failure rate of inhibitory postsynaptic currents (IPSCs) in TC neurons with effects most prominent at 30 Hz (Fig. 2B). To further explore the mechanism underlying the increased failure rate, we analyzed synaptic latency—the interval between the onset of light stimulation and the onset of IPSCs during individual events over 5 seconds. Detailed analysis revealed that CBZ increased synaptic latency at frequencies of 5, 10, and 20 Hz (Fig. 2D and Fig. S4A-B). Collectively, these findings suggest that CBZ inhibits GABAa-mediated chloride currents in TC neurons via a reduction action potential initiation and propagation in a frequency-dependent manner.

**Figure 2.**
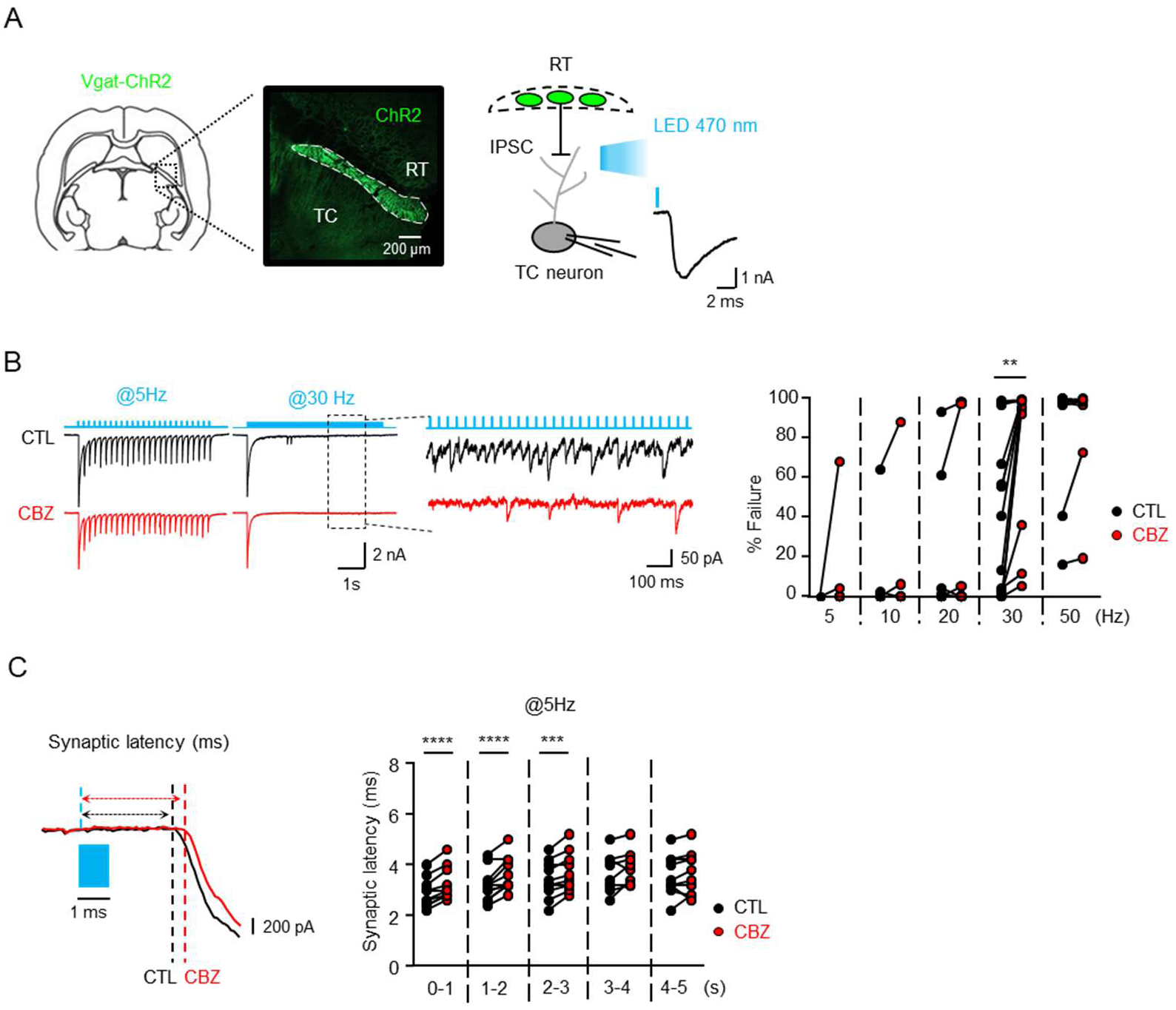
Disruption of GABAergic Transmission at the RT-TC Synapse by CBZ. (A) Representative fluorescence images (left) showing RT region expressing ChR2-YFP in VGAT-ChR2-EYFP mice, with a schematic (right) illustrating the RT-TC synaptic circuit. Optogenetic activation of RT neurons was achieved with a 1 ms blue light pulse, while 1 mM kynuric acid was applied to block glutamatergic transmission, isolating GABAergic inhibitory postsynaptic currents (IPSCs) in thalamocortical (TC) neurons. (B) Representative traces (left) showing IPSCs evoked in TC neurons by 5 Hz and 30 Hz optogenetic stimulation for 5 seconds. The graph (right) shows the percentage of synaptic failure in RT-TC inhibition across various stimulation frequencies (5, 10, 20, 30 and 50 Hz) before and after CBZ treatment (n = 12). (C) Representative traces (left) showing synaptic latency (in ms) between optogenetic stimulation onset and IPSC onset during 5 Hz stimulation and graph (right) showing quantified synaptic latency at 1-second intervals before and after CBZ treatment (n = 12). Statistical significance was determined using a paired t-test. **p < 0.01, ***p < 0.001, ****p < 0.0001.

### PV- and SOM-expressing RT neurons exhibit distinct sensitivity to CBZ at the RT-TC synapse

Considering the distinct anatomical and functional connections of the PV-and SOM-expressing RT neurons to TC neurons, as well as their diverse roles in behavior (11–13, 22), we sought to determine whether CBZ differentially affects the outputs of these two RT neuron subtypes. To address this, we introduced Cre-dependent hChR2-EYFP into the RT region of mice carrying either a PV-Cre or a SOM-Cre transgene. Robust IPSCs were observed at least three weeks post-injection, confirming reliable expression of hChR2 in RT neurons (Fig.3A). Interestingly, IPSCs at the SOM-expressing RT to TC synapse exhibited a higher failure rate compared to those at the PV-expressing RT to TC synapse during stimulation at 10, 20, and 30 Hz (Fig. 3B). CBZ further increases the failure rate at the PV-expressing RT to TC synapse during higher frequency stimulation (30 and 50 Hz; Fig. 3C). In contrast, synaptic output from SOM-expressing RT neurons was more strongly affected by CBZ, with elevated the failure rate at lower frequencies (10, 20, and 30 Hz; Fig. 3E). In addition, CBZ significantly impacted synaptic latency at both PV- and SOM-expressing RT to TC synapses during 5 Hz stimulation (Fig. 3C, 3D, 3E, and 3F). In addition, CBZ affected synaptic latency at 10 and 20 Hz, (Fig. S5A-D). Together, these findings suggest that CBZ exerts subtype—specific and frequency-dependent effects on RT to TC synaptic transmission, highlighting the differential modulation of PV- and SOM-expressing RT neuron outputs by CBZ.

**Figure 3.**
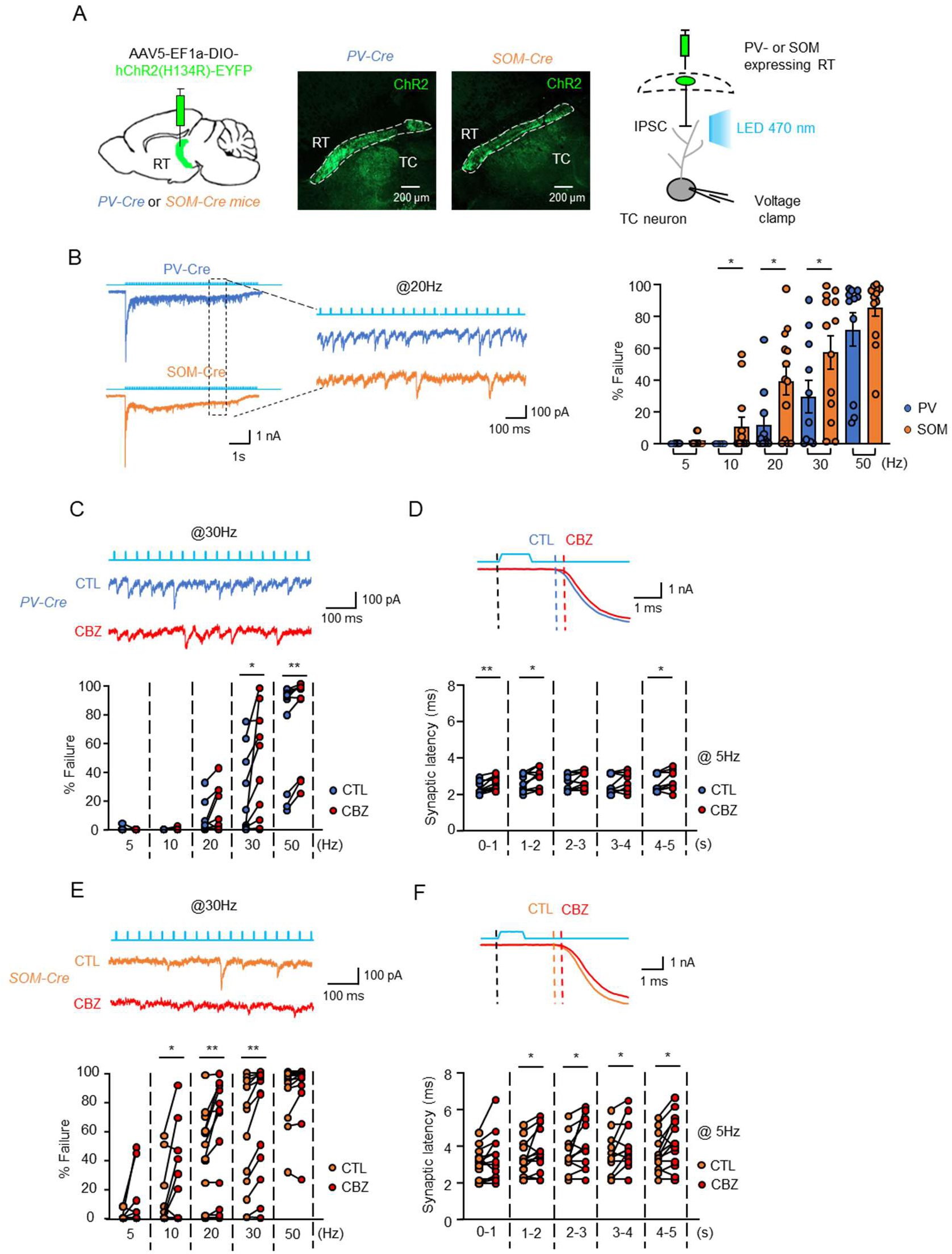
Differential Sensitivity of GABAergic Transmission at PV- and SOM-Expressing RT-TC Synapses to CBZ. (A) Schematic (left) illustrating the viral injection of AAV5-EF1a-DIO-hChR2(H134R)-EYFP into the RT region of PV-Cre and SOM-Cre mice and fluorescent image (middle) showing horizontal brain sections showing ChR2-EYFP (green) expression in RT neurons and schematic (right) representing isolated RT synaptic connections to TC neurons and optogenetic activation of ChR2-expressing RT neurons in PV-Cre and SOM-Cre mice. (B) Representative traces (left) and graph (right) showing the percentage of IPSC failures in TC neurons from PV-Cre and SOM-Cre mice during optogenetic stimulation at 5, 10, 20, 30, and 50 Hz (PV-Cre: n = 10, SOM-Cre: n = 13). (C) Representative IPSC traces (top) in PV-Cre mice during 30 Hz stimulation and graph (bottom) showing the percentage of IPSC failures at the RT-TC synapse in PV-Cre mice across different stimulation frequencies (5, 10, 20, 30, and 50 Hz) before and after CBZ treatment (n = 10). (D) Representative traces (top) of synaptic latency (ms) in PV-Cre mice during 5 Hz stimulation and graph (bottom) showing synaptic latency between optogenetic stimulation onset and IPSC onset at 1-second intervals before and after CBZ treatment (n = 10). (E) Representative IPSC traces (top) in SOM-Cre mice during 30 Hz stimulation and graph (bottom) showing the percentage of IPSC failures at the RT-TC synapse in SOM-Cre mice across different stimulation frequencies (5, 10, 20, 30, and 50 Hz) before and after CBZ treatment (n = 13). (F) Representative traces (top) showing synaptic latency (ms) in SOM-Cre mice during 5 Hz stimulation and graph (bottom) showing synaptic latency between optogenetic stimulation onset and IPSC onset at 1-second intervals before and after CBZ treatment (n = 13). Statistical significance was determined using a paired t-test. *p < 0.05, **p < 0.01.

### CBZ aggravates absence seizure and reduced locomotor activity in the absence seizure model of SCN8a^med/+^ mice

Previous report indicates that CBZ aggravates absence seizures in GAERS (5) and in a mouse model of acute absence seizures induced by γ-butyrolactone (GBL), both of which are accompanied by motor impairments including behavioral arrests (6). Based on these findings, we hypothesized that CBZ might worsen deficits in locomotor activity as well as absence seizures. To test this hypothesis, we utilized SCN8a^med/+^ mice, which exhibit spontaneous absence seizures characterized by SWD that have been previously validated in our lab (15). EEG recordings confirmed the presence of SWDs in SCN8a^med/+^ mice, a hallmark of absence seizures, which were ameliorated by ethosuximide (200 mg/kg), an anti-absence seizure medication (Fig. 4A, n = 3). In contrast, while SCN8a^WT^ mice displayed no abnormalities on EEG (Fig. S6A), administration of a low dose of pentylenetetrazol (PTZ; 20 mg/kg) effectively induced SWDs in these WT mice (Fig. S6A-B). To investigate the effects of CBZ on seizures, we administered either vehicle or CBZ (15 mg/kg, intraperitoneally) to both SCN8a^WT^ mice and SCN8a^med/+^ mice and performed EEG recordings for 1 hour. In SCN8a^med/+^ mice, CBZ significantly increases the number and total duration of SWD during the the 20 – 40 min period following injection compared to baseline (Fig. 4B-D). We further evaluated general locomotor activity using the open field test, a standard assay for measuring travel distance (Fig. 4E). During the 20-30 min period following i.p injection, SCN8a^med/+^ mice in both vehicle- and CBZ-treated groups exhibited significantly reduced travel distances compared to SCN8a^WT^ mice (Fig.4F-G). Strikingly, CBZ treatment further reduced the travel distance of SCN8a^med/+^ mice compared to the vehicle-injected groups (Fig. 4F-G) during the 30 – 40 min post-injection period (Fig.4F-G). These results demonstrate that CBZ aggravates absence seizures and reduced locomotor activity in the SCN8a^med/+^ mouse model of absence seizures, highlighting its deleterious effects on both seizure activity and motor function.

**Figure 4.**
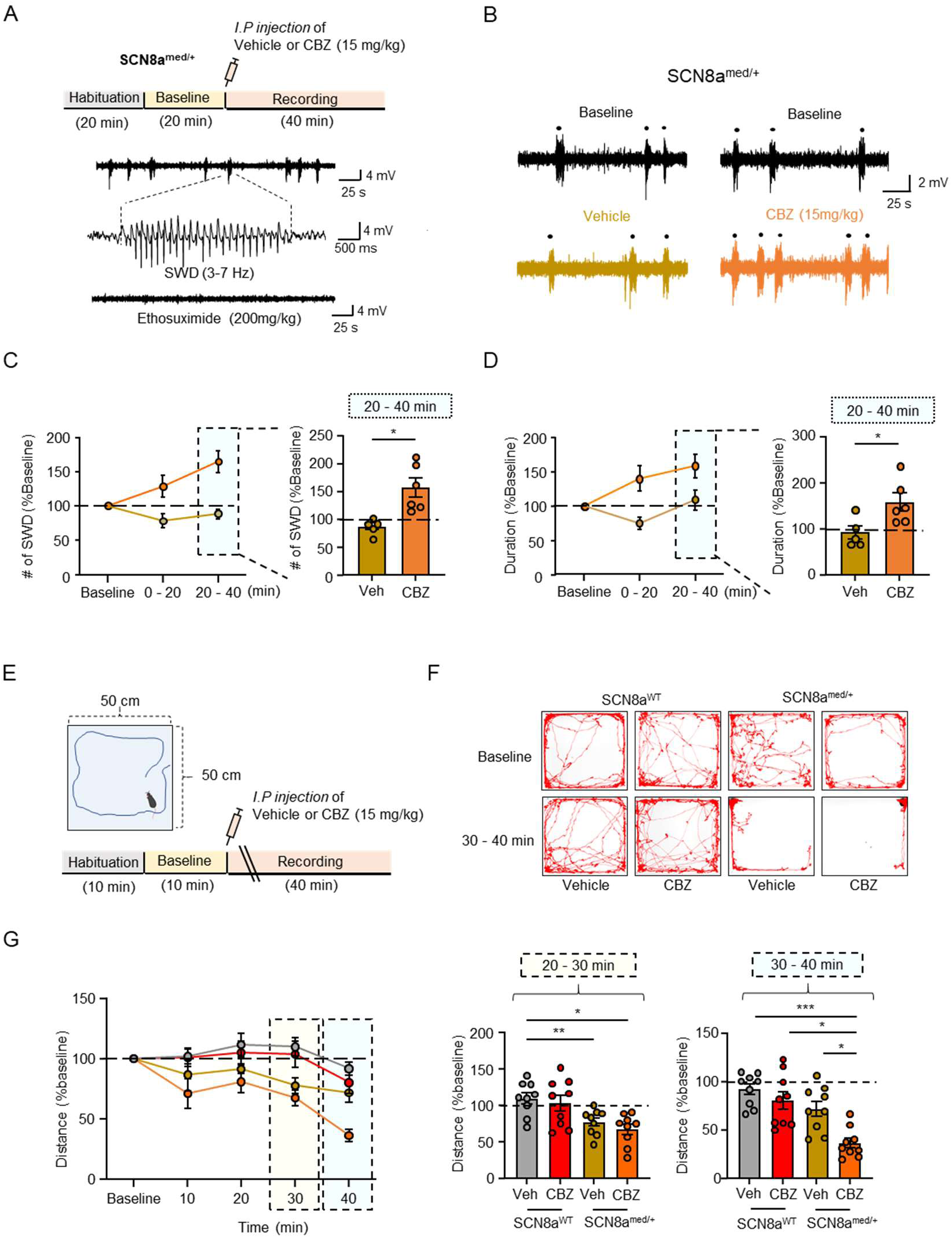
Impact of CBZ on Absence Seizures and Hypoactivity in the Absence Seizure Model of SCN8a^med/+^ Mice. (A) Experimental timeline (top) outlining the protocol for habituation (20 min), baseline recording (20 min), and treatment (40 min). Representative EEG traces (bottom) showing spike-wave discharges (SWDs, 3–7 Hz), characteristic of absence seizures, and the suppression of SWD activity following Ethosuximide administration (200 mg/kg, i.p.). (B) Representative EEG traces showing SWD activity during baseline, after vehicle, and following CBZ administration (15 mg/kg, i.p.) in SCN8a^med/+^ mice. (C) Graph (left) showing the normalized number of SWDs over a 60-minute recording period and graph (right) showing the normalized number of SWDs during the 20 - 40 minutes post-administration period following either vehicle or CBZ (15 mg/kg, i.p.) in SCN8a^med/+^ mice (Veh: n = 5, CBZ: n = 6). (D) Graph (left) showing the normalized total duration of SWDs over 60 minutes and graph (right) summarizing the normalized total duration of SWDs during the 20 - 40 minutes post-treatment period with vehicle or CBZ (15 mg/kg, i.p.) in SCN8a^med/+^ mice (Veh: n = 5, CBZ: n = 6). (E) Experimental timeline outlining the protocol for habituation (10 min), baseline recording (10 min), and post-treatment recording (40 min) in the open field. (F) Representative movement traces showing locomotor activity in the open field during baseline and following vehicle or CBZ (15 mg/kg, i.p.) administration in SCN8a^med/+^ mice. (G) Graph (left) showing the normalized distance traveled over 50 minutes during baseline and following vehicle or CBZ (15 mg/kg, i.p.) administration and graph (right) summarizing the normalized distance traveled during the 20-30 minutes and 30-40 minutes intervals post-treatment with vehicle or CBZ (15 mg/kg, i.p.) in SCN8a^WT^ (Veh: n = 9, CBZ: n = 9) and SCN8a^med/+^ mice (Veh: n = 9, CBZ: n = 9). Statistical significance was determined using a paired t-test. *p < 0.05, **p < 0.01, ***p < 0.001.

### CBZ aggravates the pre-existing reduction of RT firing in SCN8a^med/+^ mice

Previously, we reported that the reduced firing of RT neurons in SCN8a^med/+^ mice contributes to the pathogenesis of absence seizures (15). To determine whether CBZ-induced aggravation of absence seizures in SCN8a^med/+^ mice (Fig. 4C-D) results from further inhibition of tonic firing of RT neurons, we recorded APs in RT neurons using a series of square pulses (0 to 300 pA in 30 pA increments) with the same concentration of CBZ (30 µM) applied in Figure 1. Consistent with our prior findings (15), RT neurons in SCN8a^med/+^ mice exhibited a lower number of APs compared to SCN8a^WT^ mice under control conditions, particularly in response to the strongest depolarizing current injections (300 pA) (Fig. 5A). CBZ significantly reduced the number of APs during the last 500 ms, comprising tonic firing, in both SCN8a^WT^ and SCN8a^med/+^ mice (Fig. 5B and S7A). Notably, this reduction in APs was significantly greater in SCN8a^med/+^ mice compared to SCN8a^WT^ (Fig. 5B). AP properties during the first three spikes under a 240 pA current injection were minimally affected by CBZ (Fig. S8A). As observed previously (Fig. S2B), CBZ altered AP properties in SCN8a^WT^, including overshoot, threshold, amplitude, half-width, maximum and minimum dV/dt, from the last three spikes (Fig. S8B). However, these AP properties in SCN8a^med/+^ mice were not significantly affected by CBZ. (Fig. S8B). To assess the use-dependent effects of CBZ, particularly whether these effects were driven by steady depolarization during the train vs. just during the AP transient depolarization, we applied pulse train stimulation (300 pA, 5 ms per pulse) at frequencies of 5, 10, 20, 30, and 50 Hz for 5 seconds. These stimuli reliably induced single action potentials with each pulse (Figure.5C). CBZ did not affect the number of APs generated during pulse trains at any tested frequency (Fig. 5D-E). AP properties from the 1^ST^ – 3^rd^ spikes and last 3 spikes in both SCN8a^WT^ and SCN8a^med/+^ mice were not affected by CBZ (Fig. S9A-B). These findings show that CBZ aggravates the impaired of RT firing in SCN8a^med/+^ mice specifically during maintained DC step current injections. Thus, the sustained depolarization between each action potential is critical and necessary to induce use-dependent block. Activity patterns with periods of sustained depolarizations during seizures would then presumably contribute to the CBZ-induced worsening of absence seizures.

**Figure 5.**
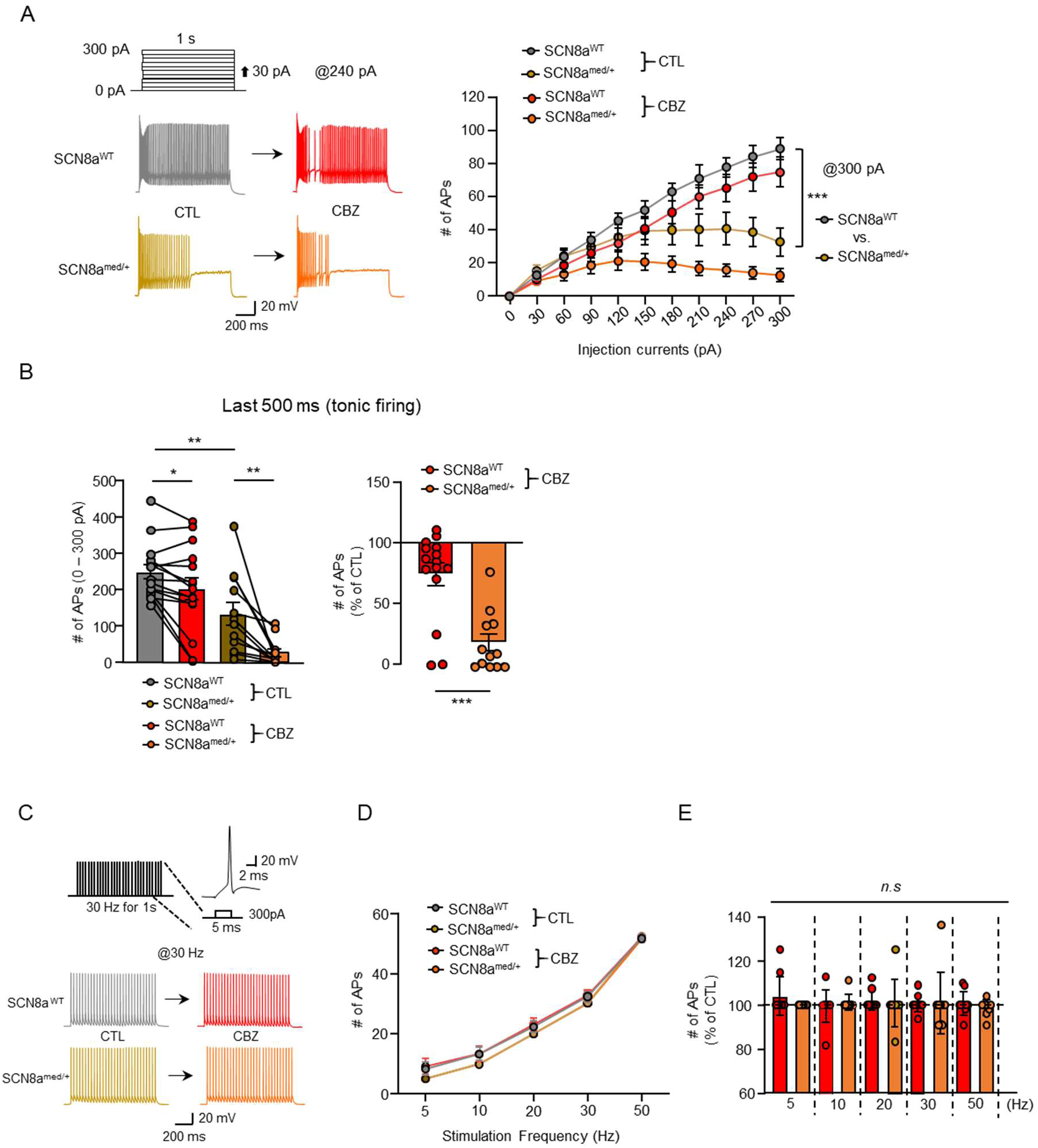
Aggravated Inhibition of Tonic Firing in RT by CBZ in SCN8a^med/+^ Mice. (A) Representative traces (left) showing the number of APs elicited by 240 pA current injection for 1-second stimulation in RT neurons before and after a 10-minute incubation with CBZ in SCN8a^WT^ and SCN8a^med/+^ mice and graph (right) depicting the number of APs elicited by current injections ranging from 0 to 300 pA under CTL and CBZ conditions in SCN8a^WT^ (n = 15) and SCN8a^med/+^ mice (n = 15). (B) Graph showing the number of APs normalized to CTL in SCN8a^WT^ and SCN8a^med/+^ mice. (C) Experimental design illustrating a pulse train (30 Hz) with 1-second 300 pA current injections (5 ms each), along with representative traces (bottom) of APs elicited by 30 Hz pulse trains before and after CBZ (30 µM) incubation in SCN8a^WT^ and SCN8a^med/+^ mice. (D) Graph showing the number of APs elicited by 5, 10, 20, 30, and 50 Hz pulse trains under CTL and CBZ conditions in SCN8a^WT^ (n = 10) and SCN8a^med/+^ mice (n = 9). (E) Graph showing the number of APs normalized to CTL in SCN8a^WT^ and SCN8a^med/+^ mice across pulse trains at different frequencies (5, 10, 20, 30, and 50 Hz). Statistical significance was determined using a paired t-test. *p < 0.05, **p < 0.01, ***p < 0.001.

### CBZ does not change the disruption of GABAergic transmission at the RT-TC synapse in SCN8a^med/+^ mice

Finally, we assessed whether CBZ further suppresses GABAergic transmission at the RT-TC synapses, which are already compromised in SCN8a^med/+^ mice. Initially, we confirmed that compared to SCN8a^WT^ mice, SCN8a^med/+^ mice exhibit an increased failure rate at 20 Hz stimulation (Fig. 6A), with little change in synaptic latency at 5 and 10 Hz stimulation (Fig. 6B and Fig. S10A). CBZ significantly increased the failure rate at 20 Hz stimulation in SCN8a^WT^ mice without further suppression in SCN8a^med/+^ mice (Fig. 6C). Consistent with previous findings (Fig. 2D), CBZ also prolonged synaptic latency at 5 and 10 Hz stimulation in both genotypes (Fig. 6D, Fig. S10B). Collectively these findings indicate that CBZ does not further exacerbate inhibitory effects at the RT-TC synapse in SCN8a^med/+^ mice.

**Figure 6.**
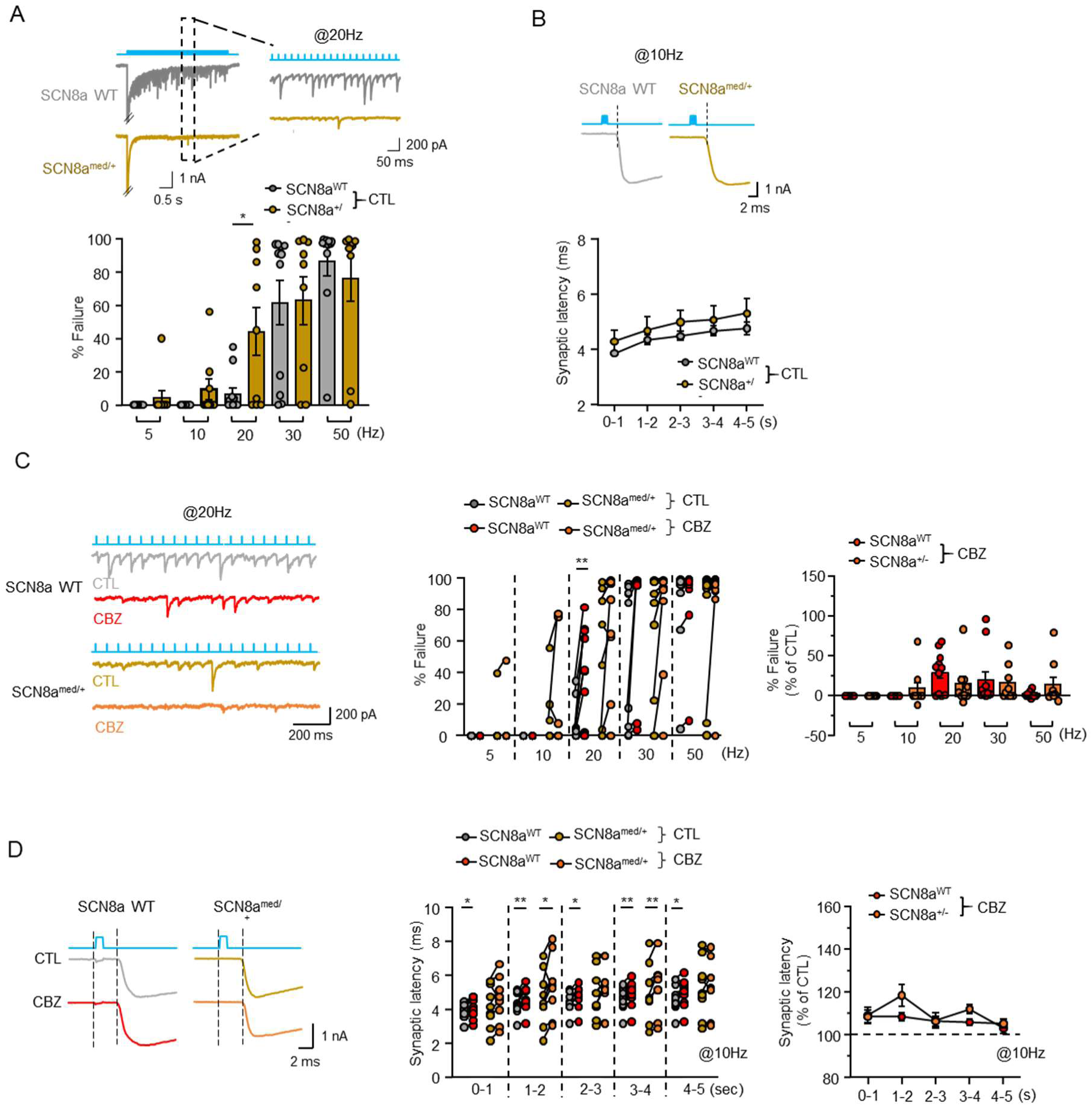
CBZ Exerts No Significant Additional Effect on GABAergic Transmission at the RT-TC Synapse Between SCN8a^WT^ and SCN8a^med/+^ Mice. (A) Representative traces (top) showing inhibitory postsynaptic currents (IPSCs) in TC cells evoked by 20 Hz optogenetic stimulation of RT regions for 5 seconds and graph (bottom) showing the percentage of failure in RT-TC synaptic transmission at varying stimulation frequencies (5, 10, 20, 30, 50 Hz) in SCN8a^WT^ (n = 11) and SCN8a^med/+^ mice (n = 9) expressing VGAT-ChR2-EYFP. (B) Representative traces (top) showing synaptic latency between optogenetic stimulation and IPSC onset and graph (bottom) showing quantified synaptic latency at 1-second intervals during 10 Hz stimulation in SCN8a^WT^ (n = 11) and SCN8a^med/+^ mice (n = 9) expressing VGAT-ChR2-EYFP. (C) Representative traces (left) showing IPSCs evoked by 20 Hz stimulation before and after CBZ incubation for 10 minutes, graph (middle) showing the percentage of synaptic transmission failure in response to 5, 10, 20, 30, and 50 Hz stimulation, and graph (right) showing normalized failure rates to baseline across these frequencies in SCN8a^WT^ (n = 11) and SCN8a^med/+^ mice (n = 9). (D) Representative traces (left) showing synaptic latency at 10 Hz stimulation before and after CBZ treatment for 10 minutes, graph (middle) showing quantified synaptic latency at 1-second intervals, and graph (right) showing normalized latency to baseline in SCN8a^WT^ (n = 11) and SCN8a^med/+^ mice (n = 9). Statistical significance was determined using a paired t-test. *p < 0.05, **p < 0.01.

## Discussion

This study elucidates the potential mechanisms behind CBZ’s paradoxical aggravation of absence seizures using electrophysiological and behavioral assessments. CBZ selectively inhibits the tonic firing mode in RT neurons in an activity-dependent manner and suppresses GABAergic transmission at the RT-TC synapse, likely altering thalamocortical oscillatory synchronization, a hallmark feature of absence seizure. In the SCN8a^med/+^ mouse model, which exhibits spontaneous absence seizure due to reduced tonic firing in RT neurons (15), CBZ aggravate the pre-existing reduction in tonic firing of RT neurons in SCN8a^med/+^ mice, accompanied by an increase in both the number and duration of SWD. These findings provide insights into the poorly understood phenomenon of CBZ-induced worsening of absence seizures, shedding light on managing epilepsy treatment.

CBZ is an effective anti-convulsant drug for adults and children with partial and secondarily generalized seizures (23, 24). In children, however, CBZ exacerbates certain seizure types, including absence, atonic, tonic, and myoclonic seizures (1, 4, 25, 26). This paradoxical effect has been explored in animal models, demonstrating that high-dose CBZ (25 mg/kg) aggravates absence seizures in γ- butyrolactone (GBL)-induced and stargazer mouse models (6). Similarly, intracerebroventricular (15 µg in 4 µl) and VB thalamic (0.75 µg in 0.2 µl) CBZ microinjections prolong seizure duration, effects that are blocked by the GABAa receptor antagonist bicuculline (5), implicating the involvement of GABA signaling pathways in CBZ-induced seizure aggravation. Given the central role of RT neurons as a GABAergic modulator in thalamic oscillatory activity (11, 14, 22), focusing the research on these GABAergic neurons enables a deeper understanding of paradoxical phenomenon. In general, it is essential to compare the therapeutic concentration ranges in humans with the responses observed at corresponding concentrations in animal models. Studies in epileptic patients reported CBZ serum concentrations ranging from approximately 13 – 51 µM/L, within the therapeutic range. (27). Based on this data, we aimed to evaluate the effects of CBZ at physiologically relevant concentrations (20, 30, and 50 µM) on RT neurons. Our findings showed that 50 µM CBZ induced a 50% reduction in APs, while 30 µM CBZ caused a moderate yet significant reduction in APs. To better understand CBZ’s blocking mechanisms in RT’s APs, we settled on 30 µM CBZ, a concentration that reliably produced detectable effects on AP generation (17, 19), yet was unlikely to be the toxic range in which side effects such as coma, respiratory failure, and cardiac conduction defects would be notable (28, 29). Given the impairment of RT excitability and GABAergic transmission at the RT-TC synapse in thalamic hypersynchrony and absence seizure (13, 15, 22, 30, 31) along with our observation that CBZ increased the number and total duration of SWD in SCN8a^med/+^ mice, we propose that CBZ’s inhibitory effect on RT excitability might underlie aggravation of absence seizure.

CBZ inhibits voltage-dependent Na^2+^ channels through two primary mechanisms (32–36): (1) blocking Na^2+^ channels in their resting state at hyperpolarized membrane potentials and (2) inducing an activity-dependent block, an action enhanced during sustained depolarization. This results in a progressive reduction in spikes during high-frequency, but not low-frequency firing, supported by findings of enhanced CBZ inhibition of sodium channels in neuroblastoma cells in response to high-frequency stimulation (2Hz or higher) (37). Given the properties of RT neurons, especially their very high-frequency firing (up to 500 Hz), these cells might be more susceptible to CBZ inhibition compared to other neuron types such as CA1 pyramidal neurons, which exhibit minimal inhibition even at high CBZ concentrations (100 µM) (16). In addition, CBZ had variable effects on the firing behavior among three classes of GABAergic interneurons such as a basket cell (BC), a proximal dendritic targeting cell (PD), and an oriens lacunosum-moleculare (OLM) interneurons at very high rates (38, 39) in the rodent hippocampus (19).

In our study, CBZ’s efficacy was maximal in response to long-lasting, several seconds long, square pulse current injections, which simulate paroxysmal depolarization shifts, mimicking the electrical activity of an epileptic focus. Notably, this inhibitory effect disappears in response to pulse train stimulations (5 – 50 Hz) producing comparable number of total action potentials, confirming that it is not high frequency firing, per se, that engages CBZ block, but that is the sustained depolarization between action potentials. Thus, an epileptic-like plateau with sustained, seconds-long, depolarization of membrane potential to values positive of -60 mV is particularly effective in triggering use-dependent block. Note that high-frequency burst firing of RT neurons (Fig. 1B) does involve AP rates of several hundred Hz, but that each burst is limited to ∼100 ms, which is insufficient to engage use-dependent block (Fig. 1E, first 3 spikes). This is consistent with our findings that steady depolarization produces much greater block (Fig. 1F). Lastly, given its use- and frequency-dependent action, CBZ might have less impact on TC neurons with lower firing frequencies. Further comparative studies and analyses of neuronal subtype susceptibility in thalamo-cortical circuit will be vital for optimizing treatment strategies.

RT neurons provide both feedforward and feedback inhibition to excitatory TC neurons, with layer 6 corticothalamic (CT) inputs inducing feedforward inhibition of TC neurons (40–43). Reciprocal connectivity between TC and RT neurons generates 7-14 Hz spindle oscillations even without cortical input (44, 45), suggesting that GABAergic output of RT to TC involves the generation of rhythmicity in the thalamus. Recent studies demonstrated that impaired thalamic GABAergic transmission in NLG2 KO mice is correlated with spontaneous SWDs and behavioral arrest (31), strongly supporting that CBZ-induced suppression of GABAergic transmission at the RT-TC synapses presumably contributes to aggravation of absence seizures. RT neurons are heterogenous, divided into two populations based on PV and SST expression, each with distinct thalamocortical projections. PV-expressing RT neurons, with rhythmogenic properties mediated by robust expression of T-type calcium channels, mainly project to the sensory circuits of VB thalamus, modulating somatosensory behavior and seizures (11), whereas SST-expressing RT neurons target the intralaminar (IL) thalamocortical nuclei and PF thalamus, influencing gamma rhythms and visual processing (12, 14). Considering the causal link between the activity of PV-expressing RT neurons and absence seizures, increased synaptic failure at PV-expressing RT-TC connections by CBZ might underlie its seizure-aggravating effects. In the thalamocortical circuit, cortical input to the sensory thalamus induces feed-forward inhibition of TC neurons via RT neurons during oscillations. Reduced cortico-RT projection strength in GluA4-deficient (Gria4^−/−^) mice disrupts the rhythmogenic cortico-thalamo- cortical system, resulting in spontaneous SWD (42). Future studies should examine how CBZ’s impact on other synapses within thalamo-cortical circuits to clarify its role in aggravating absence seizures. Furthermore, as previously proposed, intra-RT synapses (46, 47) might contribute to absence seizure under conditions where intra-RT inhibition is disrupted. For instance, an increase in inhibitory input to RT neurons is correlated with reduction in the duration and power of oscillations, highlighting the critical role of intra-RT inhibition in modulating thalamocortical network activity (48). In this work, we also observed intra-RT synaptic responses, however, technical challenges limited the stability and duration of recordings, precluding a comprehensive evaluation the effects of CBZ. Such experiments required stable recordings and responses for greater than 30 minutes to allow for wash in and wash out of drugs, and the optogenetically triggered intra-RT responses were not stable under these conditions. Nonetheless, we anticipate that CBZ’s influence on intra-RT synapses would parallel its effects on the RT-TC synapses.

In conclusion, this study elucidates mechanisms through which CBZ aggravates absence seizures, particularly through its selective inhibition of tonic firing in RT neurons and its suppression of GABAergic transmission. Our findings demonstrate that CBZ further aggravates the pre-existing reduction in tonic firing in SCN8a^med/+^ mouse, which is correlated with an increase in both the frequency and duration of seizures. This study shed light on how CBZ alters RT neuron firing and GABAergic transmission, providing a clearer understanding of its paradoxical effect on absence seizure. Future research should aim to explore the effect of CBZ on different cell types within the thalamus, contributing to rhythmicity at the thalamocortical circuits.

## Materials and Methods

### Animals

WT mice (C57BL/6J) and hemizygous VGAT-ChR2 mice (VGAT-mhChR2-YFP, Jax stock#: 014548), parvalbumin (PV)-Cre knock-in mice (PV-Cre, Jax Stock#: 008069), and somatostatin (SOM) - Cre Knock-in mice (SOM-Cre, Jax stock#: 013044) were obtained from Jackson Laboratory and bred with C57BL/6 J mice (Stock#: 000664). Mice with the heterozygous loss of function mutation in *Scn8a* were purchased from the Jackson laboratory (C3HeB/FeJ-Scn8a^med/J^, <u>Kohrman et al., 1996</u>, Stock#: 003798), referred to in this manuscript as SCN8a^med+/−^. SCN8a^med+/-^ mice were maintained on a C3HeB/FeJ background resulting in litters that were either SCN8a^med+/−^or SCN8a^WT^. For optogenetic experiments in epileptic SCN8a^med+/−^ mice (15), hemizygous VGAT-ChR2 mice were crossed with SCN8a^med+/−^ mice. All pups were genotyped at postnatal day 21 using a commercial service (Transnetyx), and only mice of the desired genotypes were selected and randomly assigned to experimental groups. All mice were maintained on a reverse 12 hr dark/light cycle, and all experiments occurred during their active cycle. Males and females were used in all experiments and were 2–6 months of age. No experimental differences due to sex were observed. Food and water were available ad libitum. All experiments were approved by the Stanford Administrative Panel on Laboratory Animal Care (APLAC, Protocol #12363) and were performed in accordance with the National Institute of Health guidelines.

### Viral constructs and injection procedures

For the optogenetic activation experiments, PV-Cre or SOM-Cre mice aged 2 - 3 months underwent stereotaxic injections with AAV5.EF1a.DIO.hChR2(H134R)-eYFP.WPRE.hGH virus under isoflurane anesthesia, with the depth of anesthesia monitored and maintained throughout the procedure. The injections were administered using a microsyringe (10 µL capacity) with a 33-gauge beveled needle (WPI).

The stereotaxic coordinates of the injections were 1.1-1.2 mm posterior to Bregma, 1.9-2.1 mm lateral to the midline, 2.8-3.0 depth from the ventral surface. At each site, 300-400 nL of concentrated virus suspension was injected at a rate of 50 nl/min. To prevent backflow, the syringe was left in place for 5 minutes after each injection. The incisions were closed using tissue adhesive (Vetbond, 3M; St.Paul, MN), and the mice were allowed to recover on a heating mat until they became awake.

### In vitro slice electrophysiology

4 weeks following injections, mice were anesthetized using pentobarbital sodium (i.p., 55mg/kg) and brains were carefully removed and placed in a cold (∼4°C) oxygenated (95% O_2_/5%CO_2_) sucrose-based slicing solution containing (in mM): 234 sucrose, 11 glucose, 26 NaHCO_3_, 2.5 KCl, 1.25 NaH_2_PO_4_, 10 MgSO_4_, and 0.5 CaCl_2_ (310 mOsm). Horizontal slices (270 µm) containing RT and TC regions were sectioned using a vibratome (VT1200, Leica) at a slicing speed set to 0.07-0.08 mm/s. Slices were incubated in an interface chamber (Brain Slice Keeper 4, AutoMate Scientific, Berkeley, California) and continuously oxygenated in warm (∼32°C) artificial cerebrospinal fluid (ACSF) containing (in mM): 10 glucose, 26 NaHCO_3_, 2.5 KCl, 1.25 NaHPO_4_, 2 MgSO_4_, 2 CaCl_2_, and 126 NaCl (290 - 300 mOsm) for an hour and then transferred to room temperature (∼21-23°C) for at least an hour prior to recording. Whole-cell patch clamp of RT neurons was conducted using borosilicate glass patch pipettes (4-6 MΩ) filled with intracellular solution containing the following in mM: 126 K-gluconate, 4 KCl, 2 Mg-ATP, 0.3 GTP-Tris, 10 phosphocreatine and the pH was adjusted to 7.4 using KOH (290 mOsm).

For current-clamp recordings of RT neurons, cells were held at a resting membrane potential of approximately –70 mV with steady intracellular current throughout the recording. Experiments were performed in the presence of Kynurenic acid (1mm) and Gabazine (SR-95531, 10 μM) to block fast excitatory and inhibitory synaptic transmission, respectively. Pipette capacitance was compensated and the bridge was balanced. Evoked action potentials (APs) were elicited by injecting 1-s-duration current steps of increasing amplitude. AP traces were further analyzed and first and second derivatives (*dV/dt* and *d^2^V/dt^2^*, respectively) were calculated using pClampfit10.7 and customized Matlab (MathWorks) code. For voltage-clamp recordings of inhibitory synaptic events that include optogenetically evoked IPSCs (oIPSCs), the pipette solution contained (in mM): 135 CsCl, 10 HEPES, 10 EGTA, 2 MgCl_2_, 5 QX-314, and pH adjusted to 7.4 with CsOH (290 mOsm). Membrane potential was held at -70 mV and signals were recorded using a Multiclamp 700A amplifier with pClamp 10.7 software (Axon Instruments, San Jose, CA), at a sampling rate of 50 kHz and low-pass filtered at 10 kHz. oIPSCs were detected using custom software (Wdetecta, JRH). oIPSCs were blocked by bath application of 20 µM Gabazine at the holding potential of – 70 mV and were not detectable at – 5 mV, the expect reversal potential of Cl^−^. We excluded data obtained from unstable recordings in which (1) the series resistance increased by >20% from an initial value, or (2) holding current changed by more than 100 pA.

### Optogenetic stimulation

Mice that had previously been injected with AAV5.EF1a.DIO.hChR2(H134R)-eYFP.WPRE.hGH were used in this experiment. Vgat-ChR2 mice and SCN8a-Vgat-ChR2 mice were euthanized, and fresh brain slices were prepared for patch clamp physiology as described above. RT neurons expressing hChR2- eYFP were activated with 475 nm light using a 1.9 mW/cm^2^ light source (CoolLED). We applied 1 ms pulses of LED blue light at 5, 10, 20 ,30, and 50 Hz for 5 s to stimulate pre-synaptic release, as previously described (15).

### Electroencephalogram (EEG)

SCN8a^WT^ or SCN8a^med+/−^ mice aged 2-6 months were used for EEG recordings. Mice were anesthetized using isoflurane for the duration of the EEG implant procedure. Under anesthesia, the skull was exposed, and small burr holes on both sides were carefully drilled at specific coordinates for electrode placement. Sterilized gold pins were located within each of the skull burr holes with the pins just touching the dura, and then glued in place. A reference electrode was placed in cerebellum. After electrode placement, mice were allowed to recover for at least 3 days in their home cages to ensure stable electrode integration. EEG recordings were conducted in a quiet, controlled environment during the active cycle of the mice. For EEG data acquisition, mice were connected to an Open Ephys/Intan headstage recording system via lightweight, and flexible cables. EEG signals were amplified, and filtered at 2 kHz. The continuous EEG signals were recorded for 20 mins as a baseline. After i.p injection of vehicle or CBZ (20 mg/kg), EEG signals were recorded for 40 mins, with the first 20 minutes of acclimatization time not included in the analysis. Recorded EEG data were manually analyzed offline using MATLAB software with customized codes.

### Open field test

SCN8a^WT^ or SCN8a^med+/−^ mice aged 2-6 months were chosen for the open field test. Prior to the test, mice were acclimated to the testing room for at least 1 hour to minimize environmental stressors. The open field test was conducted in a square arena (50 cm x 50 cm x 50 cm) with white walls. Each mouse was gently placed in the center of the open field arena and habituated for 10 mins. After injecting vehicle or CBZ (15 mg/kg) into SCN8a^WT^ or SCN8a^med+/−^ mice, the mouse was placed in the arena again. Locomotor activity was recorded for 40 mins and the data from first 20 mins were excluded from analysis to allow time for brain penetration of the CBZ. To compare locomotor activity, we used the data from 20 - 40 min. Video-tracking was employed for accurate behavioral quantification. Recorded behaviors included locomotor activity (distance traveled) and were analyzed using Python code.

### Drugs

CBZ (Sigma, Cat. No. C4024) was dissolved in 10% dimethyl sulfoxide (DMSO) for in vitro slice recording and in 40% propylene, 10% ethanol, and 50% saline for in vivo measurement. Kynurenic acid was dissolved in 10% DMSO and GABAzine (SR 95531) was dissolved in water for slice cording to block ionotropic glutamate receptor and GABAa receptor, respectively. Equivalent solvent concentrations were included in control solutions. Ethosuximide (200 mg/mg) was dissolved in 0.3% tween 80, Sigma-Aldrich, ON, CA)

### Quantification and Statistical analysis

All bar graphs indicate the mean and all error bars represent ± standard error of the mean (SEM). Statistical analyses were performed using GraphPad Prism 9 (GraphPad Software, La Jolla, CA), Excel (Microsoft), and Custom MATLAB programs (MathWorks, Natick, MA). Paired Student’s t-tests were used to compare the recording before and after the application of 30 µM CBZ. Unpaired tests were used for group comparisons of SCN8a^med+/−^ their wild-type littermates. A *p* value of <0.05 was statistically significant. Statistical values are denoted as follows: *p < 0.05, **p < 0.01, ***p < 0.001, and ****p < 0.0001.

## Acknowledgements

We thank all current lab members for their feedback during entire project. We thank Cameron Glick for managing mouse lines, laboratory equipment and supplies.

## Funding sources

This work was supported by National Institute of Neurological Disorders and Stroke (NINDS) (NS34774, NS117150).

## Author Contributions

Sung-Soo Jang: Conceptualization, Software, Formal analysis, Investigation, Writing – Original Draft, Visualization, Funding acquisition; Nicole Agranonik: Software, Validation, Formal analysis; John R Huguenard : Conceptualization, Software, Writing – Original Draft, Funding acquisition.

## Competing Interest Statement

Authors have no conflict of interest to declare

---

**Supplemental Figure 1.**
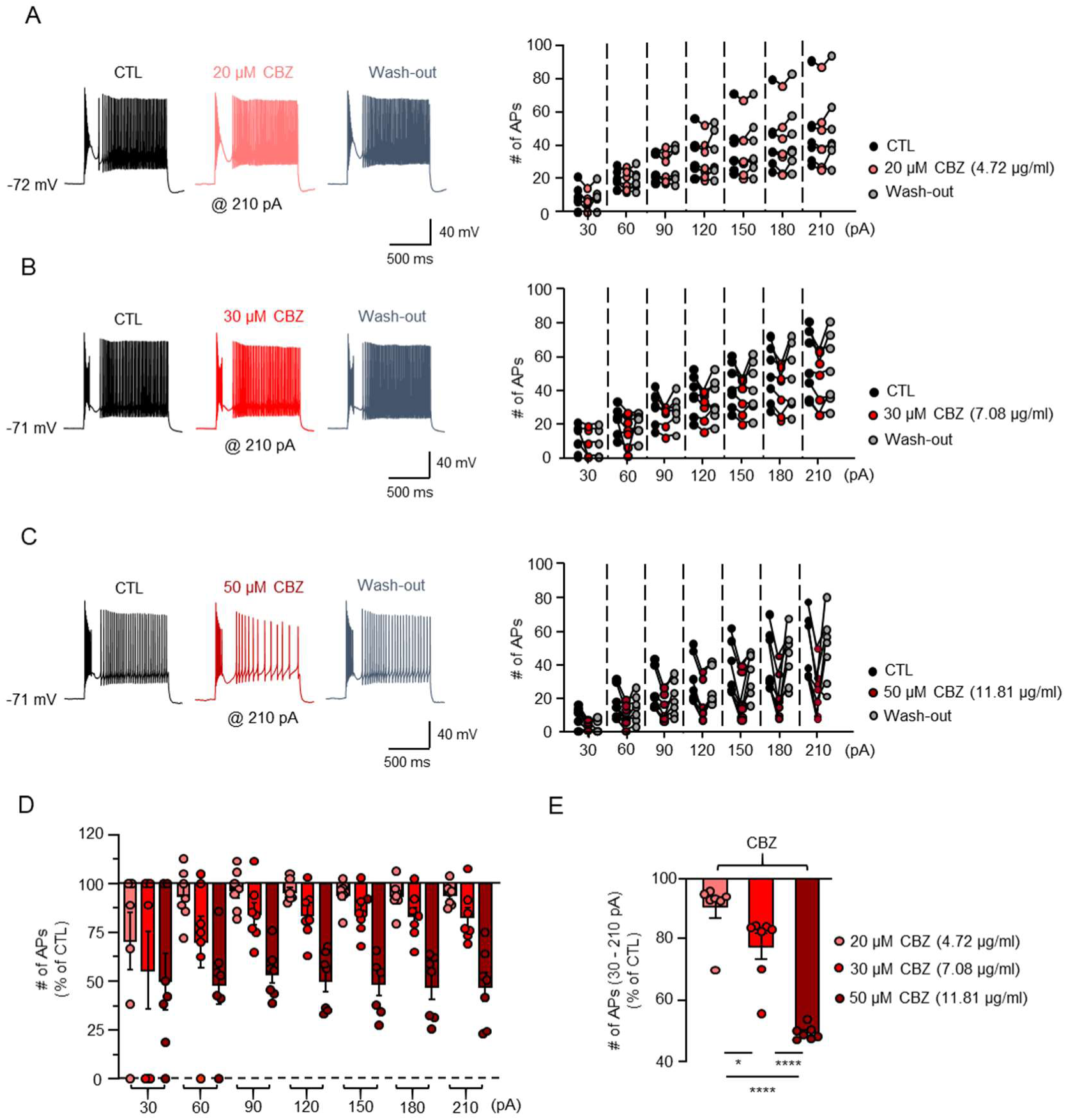
Dose-Dependent Inhibition of RT Neuronal Firing by CBZ. (A) Representative traces (left) showing AP trains elicited by 210 pA current injection from a holding potential of -72 mV and graph (right) showing the number of APs elicited by current injections (30 – 210 pA, in 30 pA increments) during 1-second stimulation epochs during CTL, 20 µM CBZ, and wash-out (n = 7). (B) Representative traces (left) showing AP trains elicited by 210 pA current injection from a holding potential of -71 mV and graph (right) showing the number of APs elicited by current injections (30 – 210 pA, in 30 pA increments) during 1-second stimulation epochs during CTL, 30 µM CBZ, and wash-out (n = 7). (C) Representative traces (left) showing AP trains elicited by 210 pA current injection from a holding potential of -71 mV and graph (right) showing the number of APs elicited by current injections (30 – 210 pA, in 30 pA increments) during 1-second stimulation epochs during CTL, 50 µM CBZ, and wash-out (n = 7). (D) Graph showing the normalized percentage reduction in the number of APs elicited by current injections ranging from 30 to 210 pA across the three CBZ concentrations (20, 30, and 50 µM). (E) Summary graph displaying the average normalized percentage reduction in the number of APs at 20 µM (n = 7), 30 µM (n = 7), and 50 µM (n = 7) CBZ concentrations. Statistical significance was determined using a paired t-test. *p < 0.05, ****p < 0.0001.

**Supplemental Figure 2.**
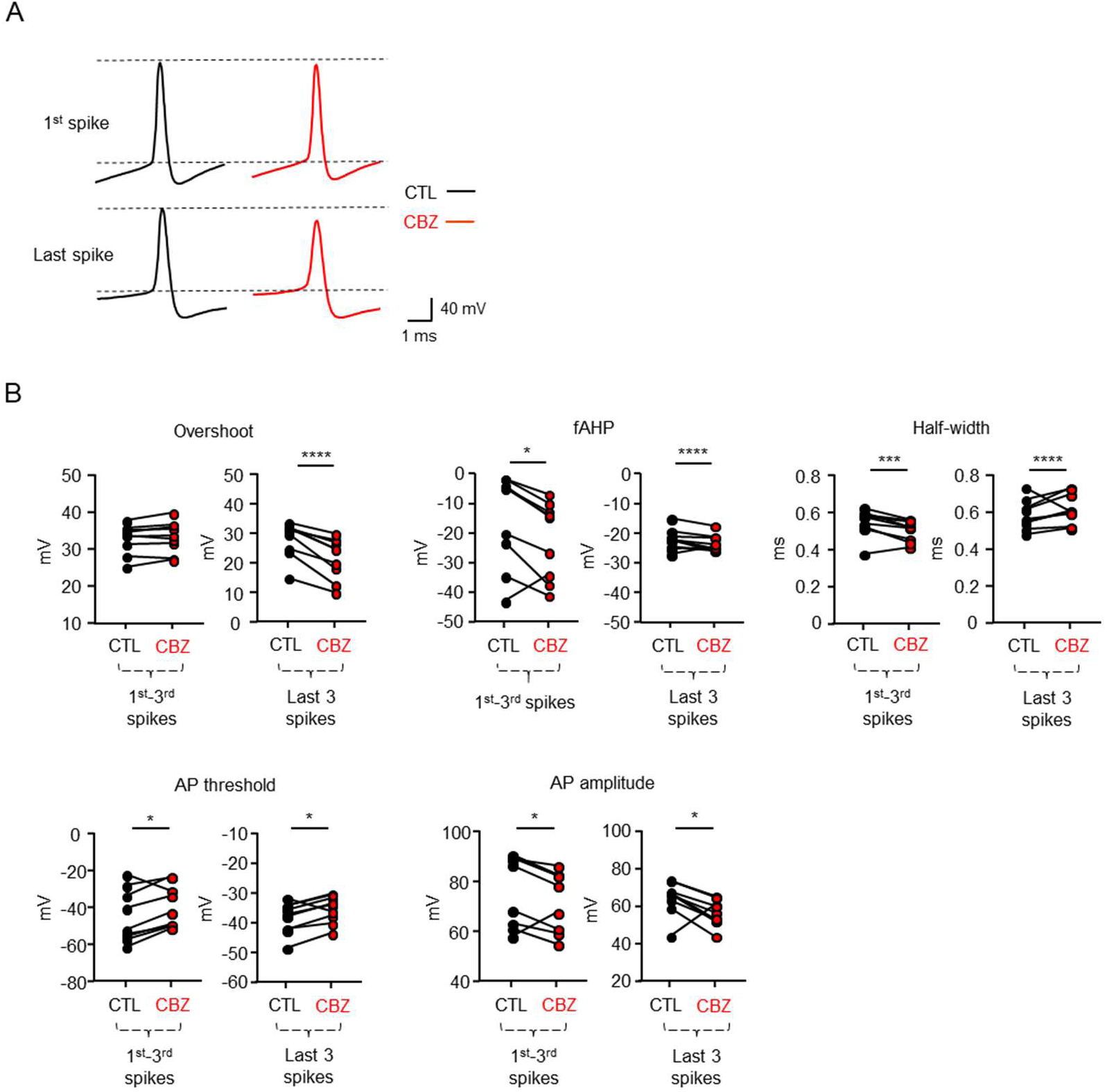
Changes in Action Potential Properties in RT Neurons Following CBZ Treatment, (A) Representative traces showing the first and last APs elicited by 300 pA current injection from a holding potential of -70 mV recorded before and after CBZ treatment. (B) Graphs displaying the average values of AP parameters, including overshoot, fast afterhyperpolarization (fAHP), half-width, AP threshold, and AP amplitude, measured from the 1st–3rd spikes and the last 3 spikes during the current injection (n = 9). Statistical significance was determined using a paired t-test. *p < 0.05, ***p < 0.001, ****p < 0.0001.

**Supplemental Figure 3.**
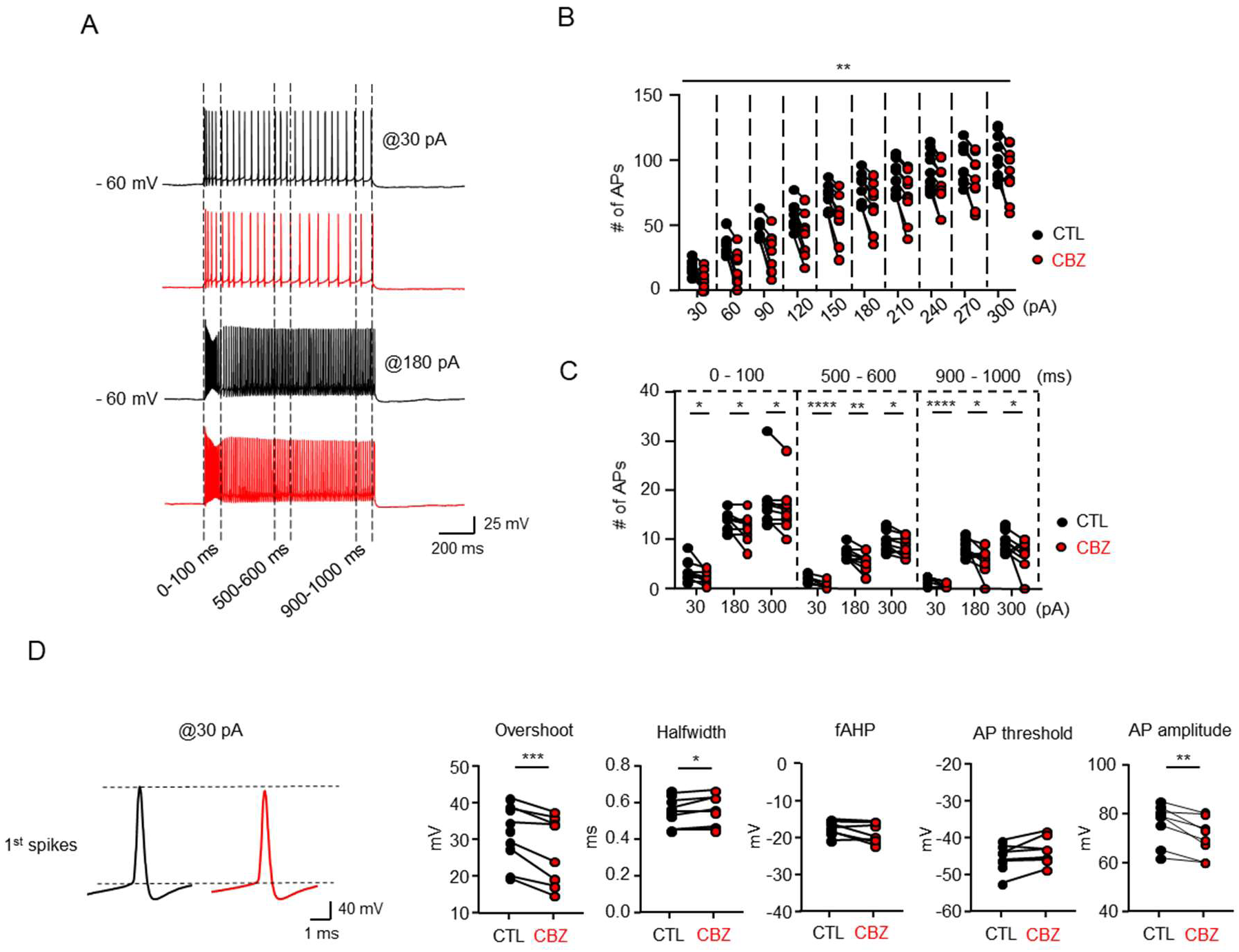
Inhibition of Tonic Firing in RT Neurons by CBZ. (A) Representative traces showing tonic firing elicited by 30 pA and 180 pA current injections from a holding potential of -60 mV, recorded before (black) and after (red) a 10-minute incubation with 30 µM CBZ (n = 9). (B) Graph showing the number of APs generated by current injections ranging from 0 to 300 pA (in 30 pA increments) during 1-second stimulation. (C) Graph showing the number of APs at specific time intervals (0–100 ms, 500–600 ms, and 900–1000 ms) elicited by current injections of 30, 180, and 300 pA during 1-second stimulation epochs. (D) Representative traces (left) showing the first AP elicited by 30 pA current injection at a holding potential of -60 mV, and graphs (right) comparing the average values of AP parameters, including overshoot, fast afterhyperpolarization (fAHP), half-width, AP threshold, and AP amplitude of the first spikes before and after CBZ treatment. Statistical analysis was performed using a paired t-test. *p < 0.05, **p < 0.01, ***p < 0.001, ****p < 0.0001.

**Supplemental Figure 4.**
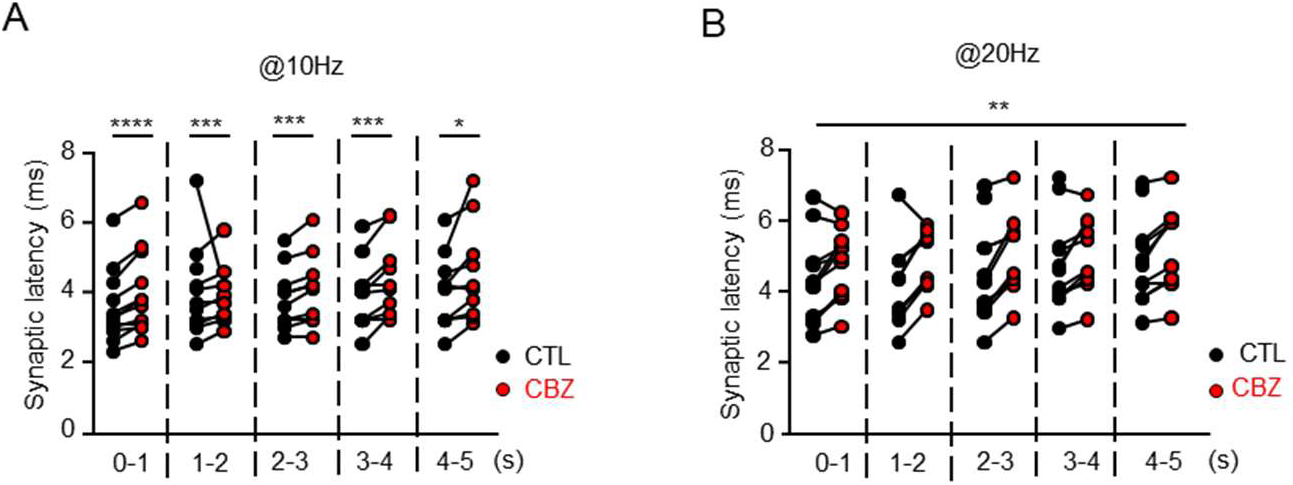
CBZ-Induced Alterations in Synaptic Latency at 10 and 20 Hz Stimulation. (A-B) Graphs showing synaptic latency measured at 1-second intervals in response to 10 (A) and 20 (B) Hz optogenetic stimulation before and after CBZ treatment (n = 12). Statistical significance was determined using a paired t-test. *p < 0.05, **p < 0.01, ***p < 0.001.

**Supplemental Figure 5.**
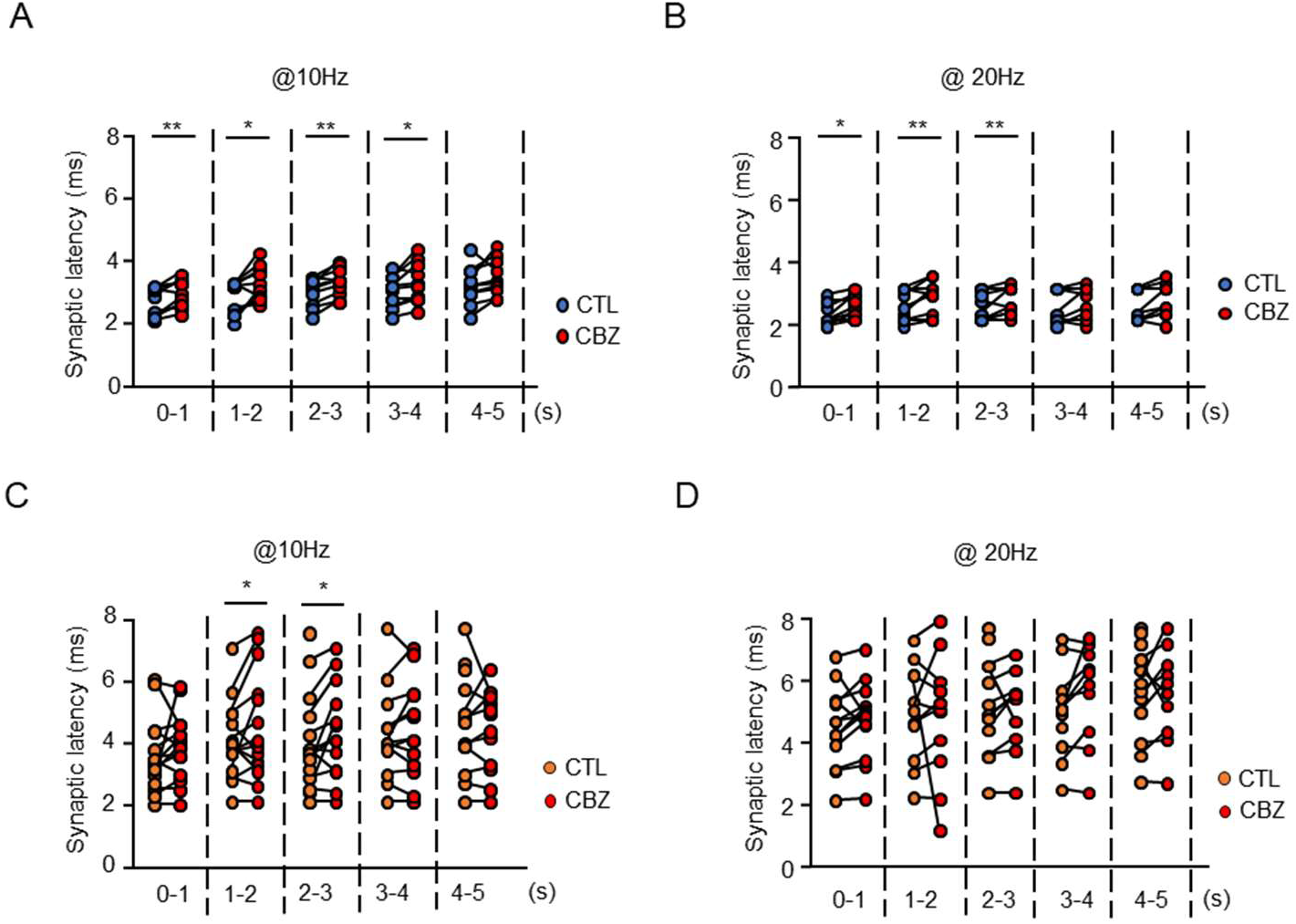
CBZ-Induced Alterations in Synaptic Latency at 10 and 20 Hz in PV-cre and SOM-cre Mice. (A-B) Graphs showing synaptic latency at 1-second intervals in response to 10 (A) and 20 (B) Hz stimulation in PV-Cre mice (n = 10). (C-D) Graphs showing synaptic latency at 1-second intervals in response to 10 (C) and 20 (D) Hz stimulation in SOM-Cre mice (n = 13). Statistical significance was determined using a paired t-test. *p < 0.05, **p < 0.01.

**Supplemental Figure 6.**
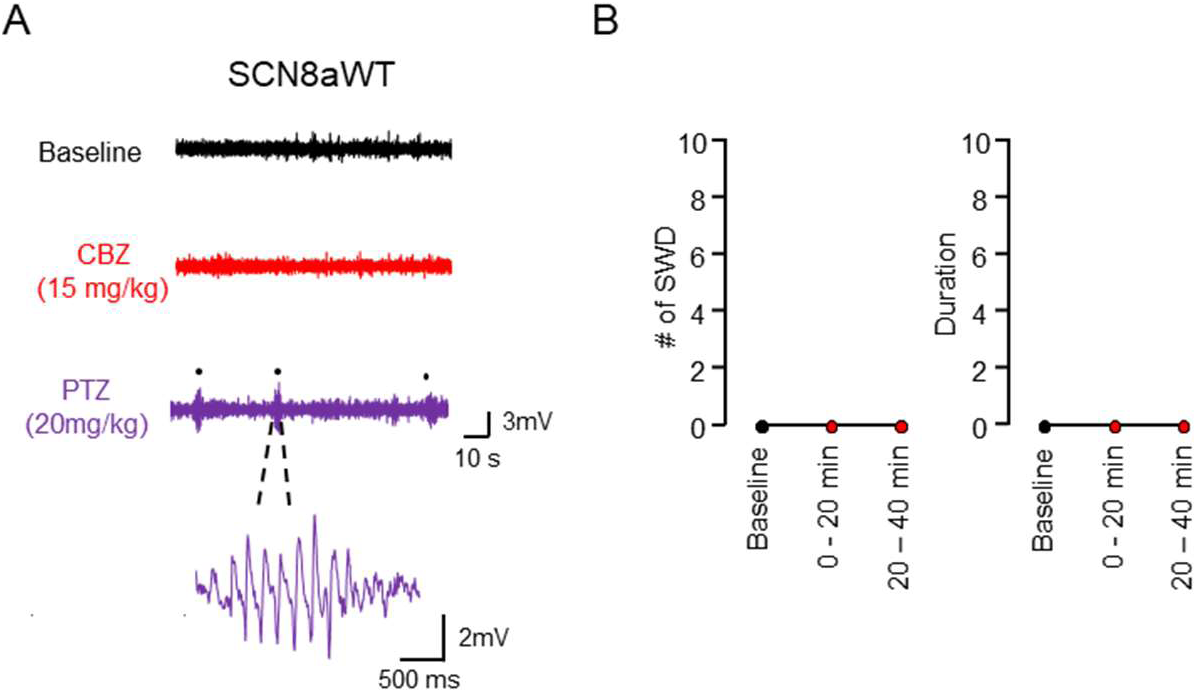
Lack of CBZ-Induced Absence Seizures in SCN8a^WT^ Mice. (A) Representative EEG traces in SCN8a^WT^ mice during baseline, after CBZ administration (15 mg/kg, i.p.), and following PTZ administration (20 mg/kg, i.p.). (B) Graphs showing the number and total duration of SWDs over a 60-minute recording period following CBZ (15 mg/kg, i.p.) administration (n = 5).

**Supplemental Figure 7.**
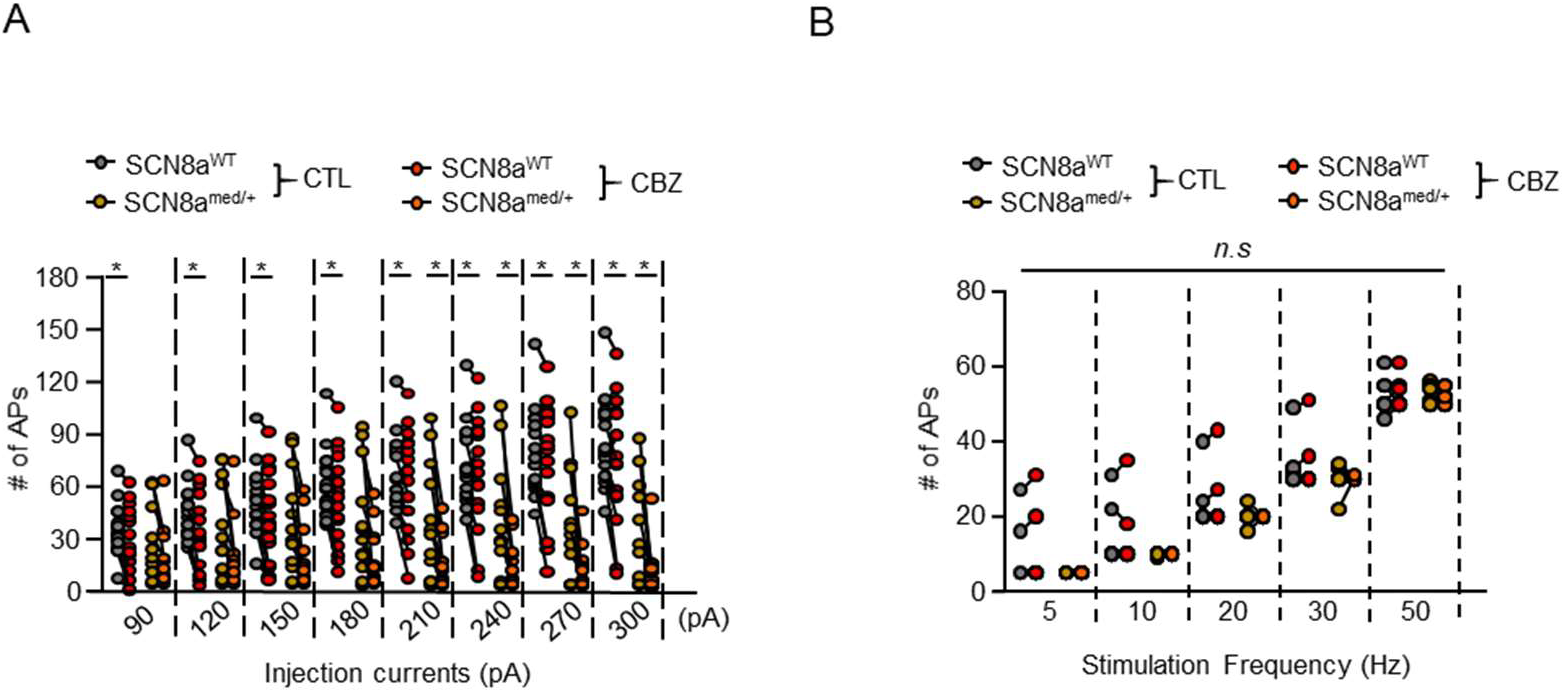
Detailed data of APs altered by CBZ in RT neurons of SCN8a^WT^ and SCN8a^med/+^ mice. (A) Graph showing the number of APs elicited by varying current injections (90 – 300 pA, in 30 pA increments) before and after CBZ in SCN8a^WT^ (n = 15) and SCN8a^med/+^ mice (n = 15). (B) Graph showing the number of APs elicited by 5, 10, 20, 30, and 50 Hz pulse trains before and after CBZ in SCN8a^WT^ (n = 10) and SCN8a^med/+^ mice (n = 9). Statistical significance was determined using a paired t-test. *p < 0.05.

**Supplemental Figure 8.**
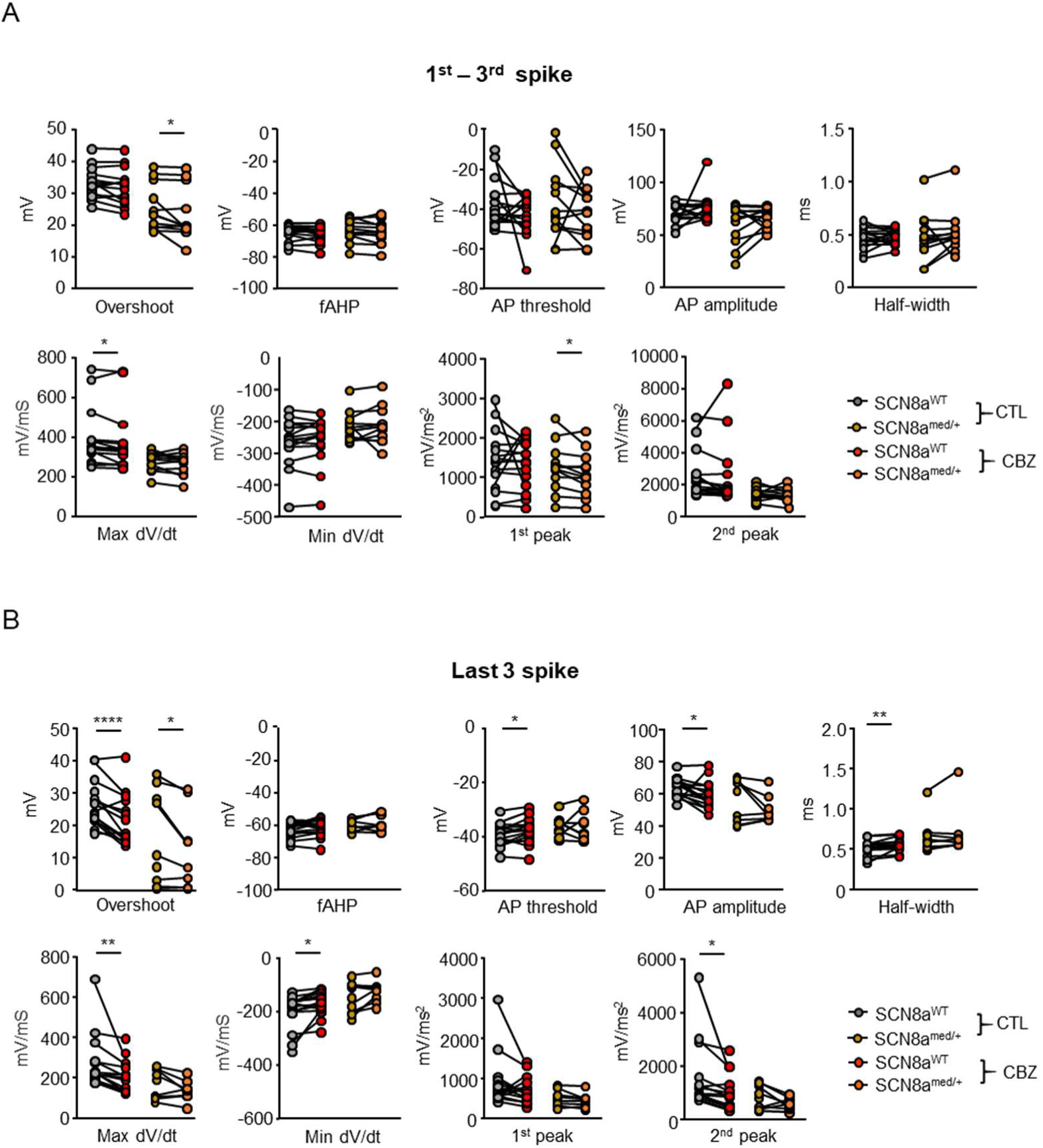
CBZ-induced changes in parameter of AP elicited by DS pulse in RT Neurons. (A) Graph showing changes in AP parameters (overshoot, fast afterhyperpolarization [fAHP], AP threshold, AP amplitude, half-width, maximum dV/dt, minimum dV/dt, and 1st/2nd peaks of d^2^V/dt^2^ for the 1st–3rd spikes elicited by 300 pA injections before and after CBZ treatment in SCN8a^WT^ (n = 15) and SCN8a^med/+^ mice (n = 15). (B) Graph showing changes in the same AP parameters for the last 3 spikes from 300 pA-elicited APs before and after CBZ treatment in SCN8a^WT^ (n = 15) and SCN8a^med/+^ mice (n = 15). Statistical significance was determined using a paired t-test. *p < 0.05, **p < 0.01, ****p < 0.0001.

**Supplemental Figure 9.**
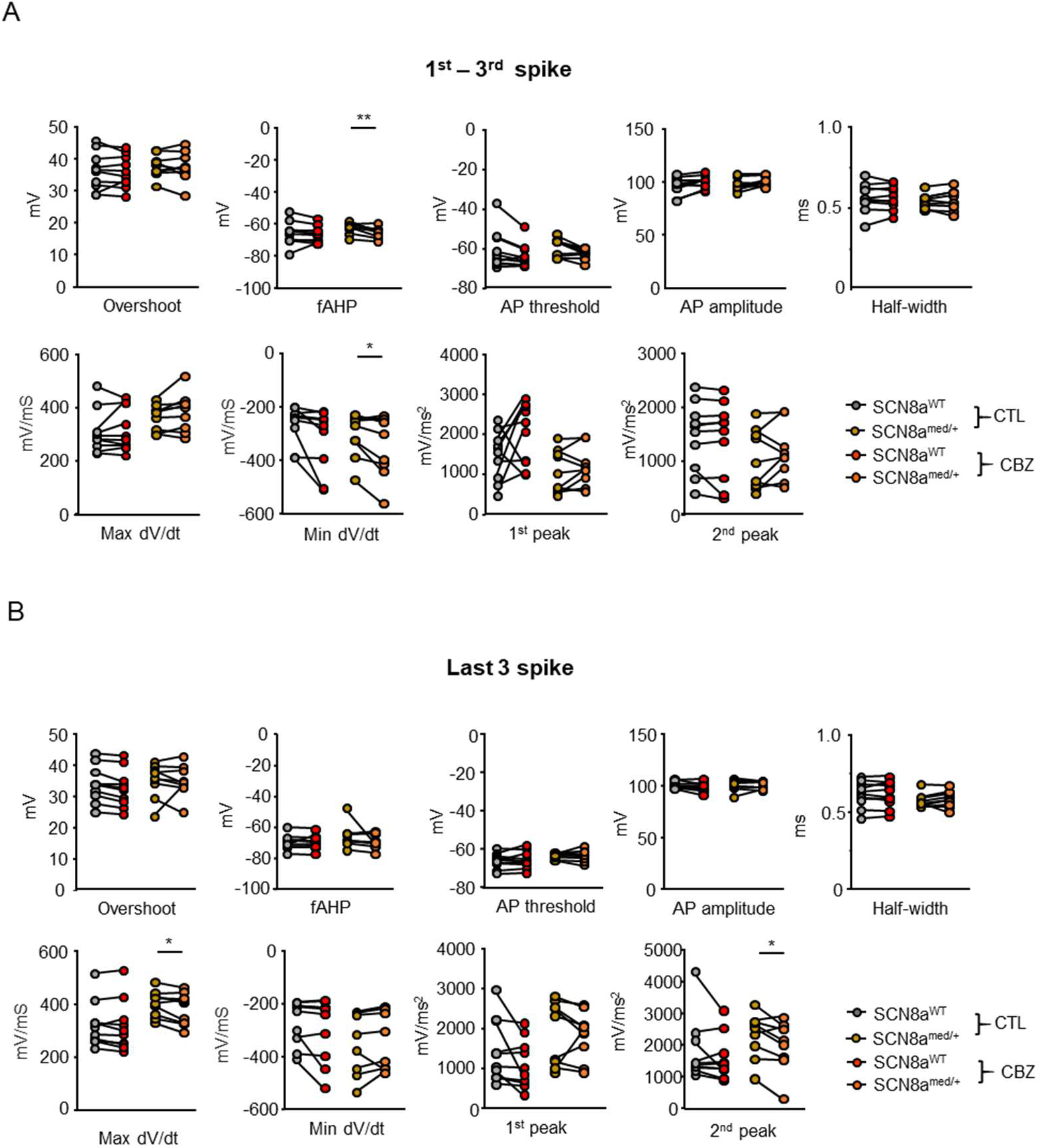
CBZ-induced changes in parameter of AP elicited by pulse trains in RT Neurons. (A) Graph showing changes in AP parameters (overshoot, fAHP, AP threshold, AP amplitude, half-width, maximum dV/dt, minimum dV/dt, and 1^st^ /2^nd^ peaks of d^2^V/dt^2^) for the 1^st^–3^rd^ spikes elicited by 50 Hz pulse trains before and after CBZ treatment in SCN8a^WT^ (n = 10) and SCN8a^med/+^ mice (n = 9). (B) Graph showing changes in the same parameters for the last 3 spikes elicited by 50 Hz pulse trains before and after CBZ treatment in SCN8a^WT^ (n = 10) and SCN8a^med/+^ mice (n = 9). Statistical significance was determined using a paired t-test. *p < 0.05, **p < 0.01.

**Supplemental Figure 10.**
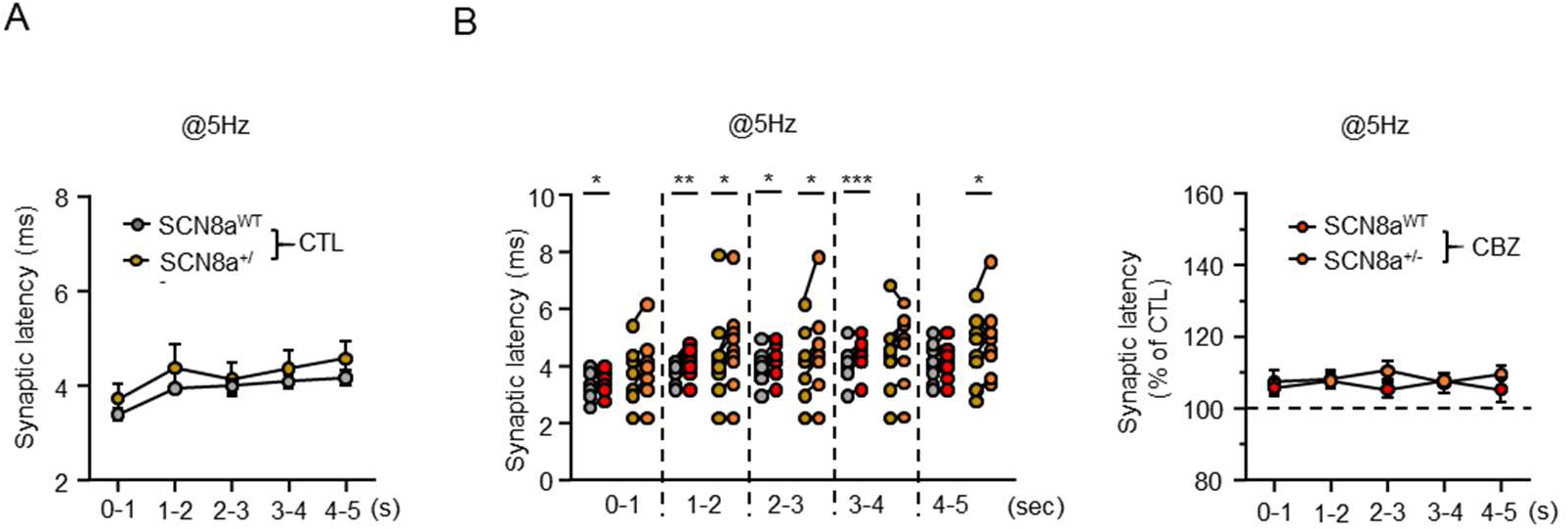
CBZ Effects on GABAergic Transmission Between SCN8a^WT^ and SCN8a^med/+^ Mice During 5 Hz Stimulation. (A) Graph showing quantified synaptic latency at 1-second intervals before and after CBZ incubation in SCN8a^WT^ (n = 11) and SCN8a^med/+^ mice (n = 9). (B) Graph (left) showing synaptic latency at 1-second intervals and graph (right) showing normalized latency to baseline in response to 5 Hz stimulation in SCN8a^WT^ (n = 11) and SCN8a^med/+^ mice (n = 9) expressing VGAT-ChR2-EYFP. Statistical significance was determined using a paired t-test. *p < 0.05, **p < 0.01, ***p < 0.001.

## Notes

### Competing Interest Statement

The authors have declared no competing interest.

